# DECIPHER integrates disentangled representation learning and prototype-based cell-type deconvolution across molecular modalities

**DOI:** 10.64898/2026.09.24.753940

**Authors:** Wenpu Lai, Chenyang Li, Qin Deng, Yue Zhu, Chenyu Liu, Zhenhua Li, Oscar Junhong Luo

**Affiliations:** Department of Systems Biomedical Sciences, School of Medicine, Jinan University, Guangzhou 510632, China

**Keywords:** Cell-type deconvolution, End-to-end learning, Bulk omics, Representation learning, Single-cell reference

## Abstract

The growing availability of single-cell resources creates new opportunities to extract cell-type-resolved information from the vast body of existing bulk omics data through cell-type deconvolution. Conventional deconvolution methods often rely on linear mixture models or specific probabilistic assumptions and can be sensitive to batch effects, whereas many deep-learning approaches are modality-specific or lack a unified end-to-end learning framework. Here, we developed DECIPHER, an end-to-end representation-learning framework for cell-type deconvolution that can be applied across multiple molecular modalities. DECIPHER learns a domain-constant representation (*Z*_c_) for deconvolution and a domain-specific representation (*Z*_s_) to model domain-associated variation. By integrating nonlinear representation learning with differentiable non-negative least-squares optimization, DECIPHER estimates cell-type proportions from *Z*_c_. Across simulated datasets, experimentally generated bulk-cell mixtures, real-world datasets, and multiple molecular modalities, DECIPHER showed robust and competitive cell-type deconvolution performance. Beyond cell-type proportion estimation, the learned *Z*_c_ supported chronological age prediction across independent cohorts and prognostic stratification in lung adenocarcinoma, demonstrating that DECIPHER can transform high-dimensional bulk omics data into low-dimensional, biologically informative representations. DECIPHER thus broadens the utility of existing bulk omics resources for biological discovery and clinical research.

## Introduction

Bulk omics profiling has long been an important approach for investigating complex biological systems, disease mechanisms, and population-level variation^1,2^. In particular, bulk-cell RNA sequencing (RNA-seq) has generated extensive transcriptomic datasets, while similar profiling strategies have also been widely applied to proteomic, metabolomic, and other molecular modalities^1,3^. Many existing bulk-cell datasets are accompanied by rich phenotypic and clinical information, including age, disease status, and treatment response, making them valuable resources for both basic and translational research. However, bulk-cell measurements represent composite molecular signals derived from multiple cell types and cellular states and therefore lack cellular-level resolution, limiting the ability to resolve sample heterogeneity and identify its underlying cellular origins^3–5^. With the rapid development of single-cell technologies, particularly single-cell RNA sequencing (scRNA-seq), and the increasing availability of comprehensive cellular atlases^6,7^, increasingly refined information on cell types and cellular states has provided a foundation for resolving cellular composition from bulk data and has further accelerated the development of cell-type deconvolution methods^8,9^.

A variety of computational strategies have been developed to infer cellular composition from bulk omics data. Conventional statistical approaches typically model bulk profiles as mixtures of cell-type-specific signals^10–12^. For example, MuSiC^8^ improves cell-proportion estimation by leveraging cross-individual expression variability to identify stable cell-type features, thereby providing an interpretable framework for deconvolving complex tissues. However, their reliance on predefined mixture assumptions may limit their ability to capture nonlinear relationships between reference and target datasets^3^. Deep-learning approaches offer more flexible nonlinear mappings: Scaden^13^ uses neural networks to directly infer cell-type proportions from bulk-cell expression profiles, TAPE^14^ incorporates tissue-adaptive strategies to improve generalization to real bulk samples, and DECODE^15^ integrates representation learning, domain adaptation, and denoising within a multistage framework for deconvolution across molecular modalities. Beyond transcriptomic deconvolution, specialized methods have further extended cellular composition analysis to additional molecular modalities and spatial contexts^3^. For proteomic data, scpDeconv^16^ addresses distributional differences between single-cell and bulk-cell measurements to estimate cellular composition. In spatial transcriptomics, Cell2location^17^, Tangram^18^, and CARD^19^ employ probabilistic modeling, cross-dataset mapping, and spatial constraints, respectively, to resolve spatially organized cellular mixtures. Together, these developments have substantially broadened the scope of cell-type deconvolution^19^.

Despite these advances, many existing methods primarily focus on estimating cell-type proportions, which alone cannot fully summarize the biological information contained in heterogeneous bulk samples^20^. Complex phenotypes such as age, disease status, and treatment response may involve molecular variation beyond changes in cellular composition^21,22^, motivating the learning of sample-level representations that support both deconvolution and downstream phenotype analyses^23,24^. Learning such representations must also account for distributional differences between reference-derived training data and real target samples, which may reflect both technical effects and biological variation^14,16,25^. Evidence from single-cell data integration suggests that excessive domain alignment may compromise the preservation of biological structure^26^. These considerations highlight the need to model domain-associated variation while retaining information relevant to cellular composition and downstream biological analysis^27^.

To address these challenges, we developed DECIPHER (DECoupled Invariant Prototypes for Heterogeneous Expression Reconstruction), an end-to-end representation-learning framework for cell-type deconvolution that can be applied across multiple molecular modalities. Using single-cell reference data to generate simulated mixtures ^13,14^, and construct cell-type prototypes, DECIPHER jointly learns from mixtures with known compositions and unlabeled target samples. At its core, DECIPHER learns two complementary latent representations: a domain-constant representation (*Z*_c_) capturing biologically relevant signals for cell-type proportion estimation while preserving information shared across data domains, and a domain-specific representation (*Z*_s_) capturing domain-associated variation, including batch effects and platform differences. By integrating nonlinear representation learning with differentiable non-negative least-squares (NNLS) optimization, DECIPHER estimates cell-type proportions from *Z*_c_ using the cell-type prototypes for multi-omics. We systematically evaluated DECIPHER in separate reference-target settings spanning transcriptomic, proteomic, metabolomic, spatial transcriptomic, and regulatory-genomic applications. These evaluations used simulated datasets, experimentally generated bulk-cell mixtures, and real-world datasets to assess deconvolution accuracy, generalization, and robustness to data perturbations and incomplete references. Beyond cell-type deconvolution, downstream phenotype analyses further demonstrated that *Z*_c_ retained biologically informative sample-level signals that supported immune-age prediction across independent peripheral blood mononuclear cell (PBMC) cohorts and prognostic stratification in lung adenocarcinoma. Together, these results establish DECIPHER as a unified framework that connects interpretable cellular composition analysis with biologically informative sample representation learning across heterogeneous molecular data.

## Results

### Overview of the DECIPHER Framework

We developed DECIPHER as an end-to-end deep learning framework that jointly performs cell-type deconvolution and latent representation learning within a unified optimization process (Fig. 1A; Methods). DECIPHER first generates simulated bulk omics (e.g., RNA-seq) profiles with known cellular compositions from single-cell reference data (e.g., scRNA-seq) and jointly models them with real bulk omics profiles. For each set of input data, including data of both simulated and real samples, the encoder disentangles the input information into two latent representations: a domain-constant representation *Z*_c_ and a domain-specific representation *Z*_s_. *Z*_c_ primarily retains biological information shared across data domains (i.e., simulated or real), whereas *Z*_s_ is explicitly guided by a domain information (e.g., dataset-specific source and generation protocol) prediction task to capture batch effects and possible additional noise and signals from unknown cell types (Methods). For the *Z*_c_ latent space, DECIPHER estimates cell-type proportions through a differentiable NNLS solver, where the *Z*_c_ representation of each sample is modeled as a non-negative combination of the cell-type latent prototypes *Z*_ct_ (Fig. 1A-B; Methods). Meanwhile, the model aligns the *Z*_c_ representations of simulated bulk and real samples to promote consistency of shared biological information across data domains. In contrast, *Z*_s_ is guided by the domain prediction task to retain domain-specific variation, particularly batch-associated information. The decoder further reconstructs the input expression profiles using both *Z*_c_ and *Z*_s_, providing a constraint that preserves informative features from the input data in the learned latent representations (Fig. 1A).

**Fig. 1:**
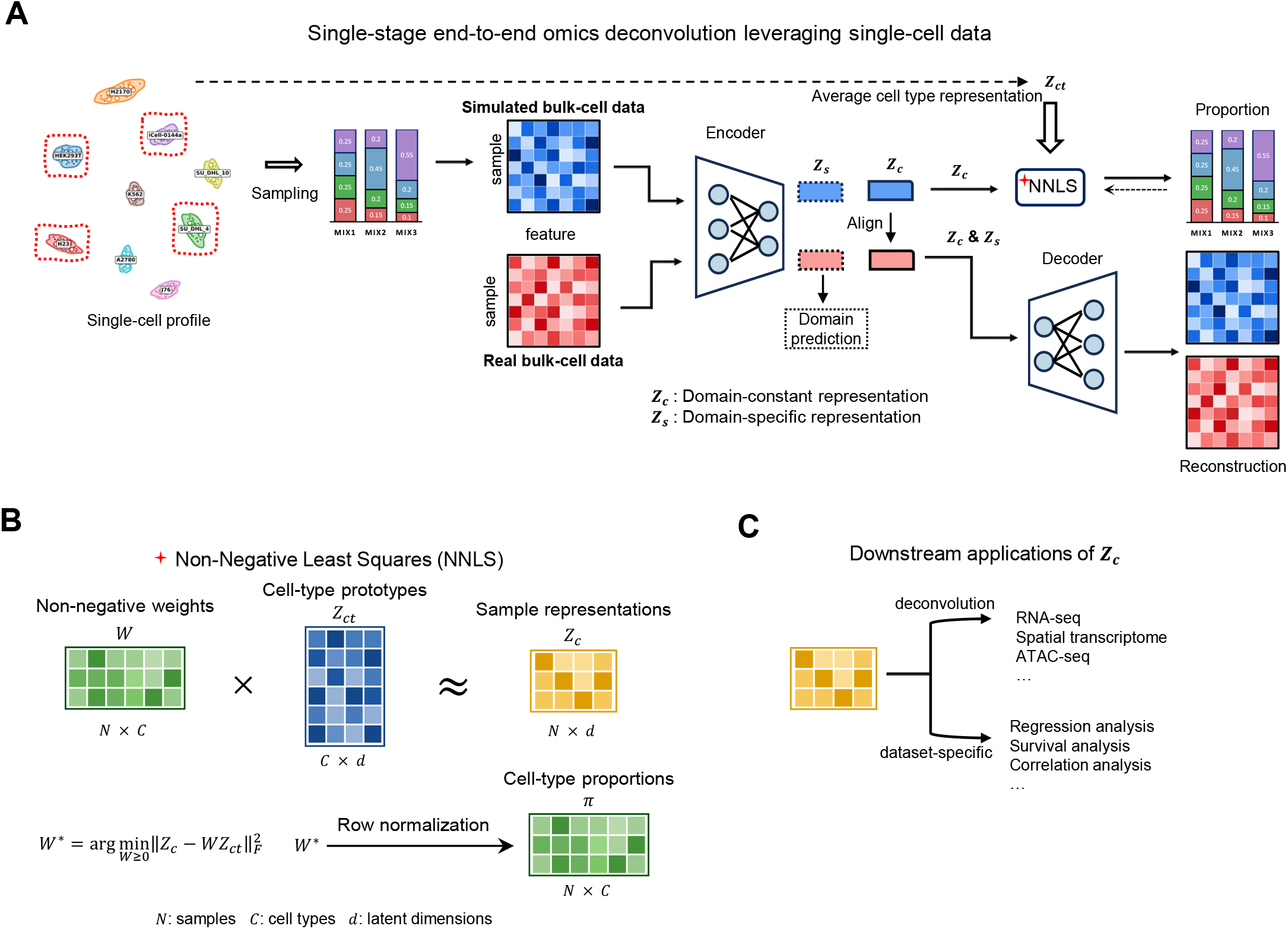
Overview of the DECIPHER framework and its downstream applications. (A) Schematic overview of DECIPHER, a single-stage, end-to-end framework for cell-type deconvolution from omics data. Single-cell transcriptomic profiles are sampled to generate simulated bulk omics profiles, which are jointly modeled with real bulk profiles. The encoder decomposes each sample into a domain-constant representation (*Z*_c_) and a domain-specific representation (*Z*_s_). Domain alignment minimizes the distributional discrepancy between the *Z*_c_ representations of simulated and real bulk samples, whereas *Z*_s_ preserves dataset-specific variation, including batch-associated effects. Cell-type proportions are inferred from *Z*_c_ using non-negative least squares (NNLS), with cell-type latent prototypes (*Z*_ct_) used as the reference. The decoder then combines *Z*_c_ and *Z*_s_ to reconstruct the input expression profiles of both simulated and real bulk samples. (B) Latent-space estimation of cell-type proportions. The cell-type proportion vector (*π*) is estimated by NNLS under a non-negativity constraint. Specifically, NNLS determines the optimal linear combination of the cell-type latent prototypes (*Z*_ct_) that best approximates the domain-constant representation (*Z*_c_). (C) Downstream applications enabled by the learned domain-constant representation (*Z*_c_).

DECIPHER jointly optimizes cell-type proportion supervision, profile reconstruction, and latent representation learning, enabling the model to learn latent representations that preserve shared biological information across datasets. The learned *Z*_c_ representation can be used for cell-type proportion estimation of multi omics data and can further serve as a latent feature space for downstream phenotype modeling tasks, such as immune age prediction (regression analysis) and survival analysis from bulk-cell transcriptome profiles (Fig. 1C).

### DECIPHER achieves accurate and robust deconvolution across diverse molecular modalities

To systematically evaluate the generalizability of DECIPHER across diverse molecular modalities and biological contexts, we benchmarked DECIPHER against state-of-the-art deconvolution methods, including DECODE^15^, MuSiC^8^, TAPE^14^, SCADEN^13^, CARD^19^, Cell2location^17^, Tangram^18^, and scpDeconv^16^, using simulated datasets (Fig. 2A-F; Methods). Performance was evaluated using the concordance correlation coefficient (CCC)^28^, root mean squared error (RMSE), and Pearson correlation coefficient (Pearson’s r, PCC) (Methods). The benchmark datasets covered diverse molecular modalities, including transcriptomic, proteomic, and metabolomic profiles. These included transcriptomic (RNA) datasets from breast tumours^29^, pancreatic islets^30^, and PBMCs (by CITE-seq^31^: cellular indexing of transcriptomes and epitopes by sequencing), together with PBMCs CITE-seq ADT (antibody-derived tag sequencing)^32^, bone marrow metabolomic^33,34^, and breast proteomic datasets^34^. The breast tumor dataset was used to evaluate cross-subtype generalization, with models trained on estrogen receptor-positive (ER+) samples and tested on estrogen receptor-negative (ER−) samples. The pancreatic islet dataset was used to evaluate cross-dataset generalization. The PBMC CITE-seq RNA and ADT datasets provided matched transcriptomic and surface protein measurements from the same cellular system, enabling evaluation of cellular composition inference across molecular modalities. The bone marrow metabolomic dataset was used to evaluate deconvolution performance at the metabolomic level. Lastly, the breast proteomic dataset was used to assess cross-physiological-state generalization by training on premenopausal samples and testing on postmenopausal samples (Methods).

**Fig. 2:**
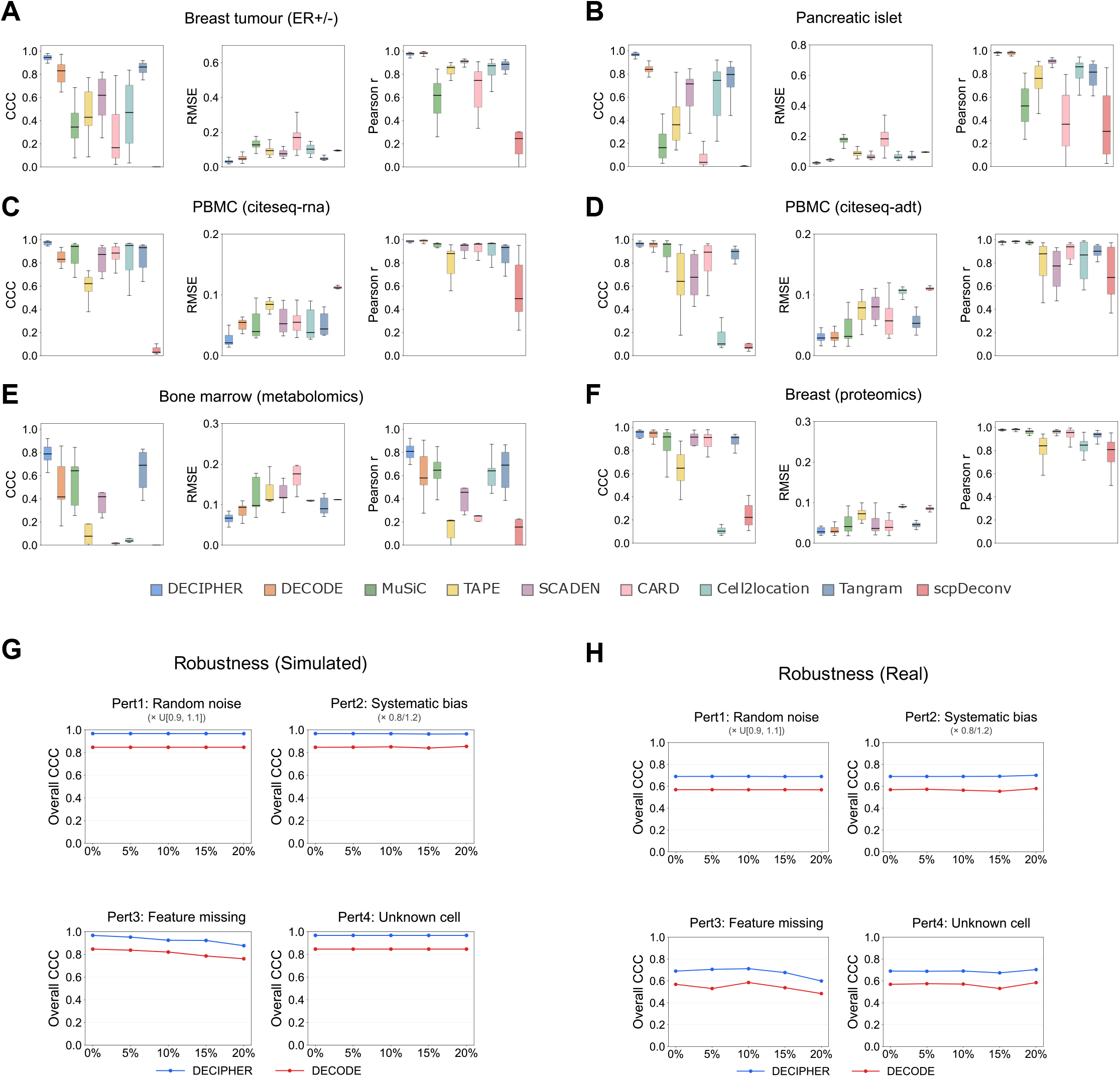
Deconvolution performance of DECIPHER across simulated bulk omics profiles and robustness to data perturbations. (A-F) Benchmarking of DECIPHER against DECODE, MuSiC, TAPE, SCADEN, CARD, Cell2location, Tangram, and scpDeconv on the breast tumor RNA-seq (A), pancreatic islet RNA-seq (B), PBMC CITE-seq RNA (C), PBMC CITE-seq ADT (antibody-derived tag) (D), bone marrow metabolomics (E) and breast proteomics (F) datasets, respectively. Performance was evaluated using the concordance correlation coefficient (CCC; higher is better), root mean squared error (RMSE; lower is better), and Pearson’s correlation coefficient (r) (higher is better). CCC and Pearson correlation range from −1 to 1, with values closer to 1 indicating better performance, whereas lower RMSE indicates better performance. Boxplots show the median (center line), 25th–75th percentiles (box), and minimum and maximum values (whiskers). (G) Robustness evaluation of DECIPHER and DECODE on simulated PBMC CITE-seq RNA-derived bulk-cell data under increasing levels of four types of perturbations: random noise, systematic bias, feature missingness, and unknown cell types. Robustness was quantified using overall CCC, calculated by pooling predicted and ground-truth cell-type proportions across all samples and cell types. (H) Robustness evaluation of DECIPHER and DECODE on real PBMC bulk-cell transcriptomic datasets from the Monaco, Microarray, and SDY67 cohorts under increasing levels of the same four perturbations. Robustness was quantified using overall CCC.

In cross-subtype and cross-dataset generalization tasks, DECIPHER maintained strong performance. On the breast tumor transcriptomic dataset, DECIPHER achieved a CCC of 0.917, an RMSE of 0.034, and a PCC of 0.966 on average (Fig. 2A), showing competitive performance compared with other approaches. On the pancreatic islet transcriptomic dataset, DECIPHER achieved a CCC of 0.956, an RMSE of 0.026, and a PCC of 0.980 on average (Fig. 2B), demonstrating stable deconvolution performance across independent datasets and experimental settings. On the PBMC CITE-seq RNA dataset, DECIPHER achieved on average a CCC of 0.967, an RMSE of 0.026, and a PCC of 0.986 (Fig. 2C), while on the matched PBMC CITE-seq ADT dataset, it achieved on average a CCC of 0.929, an RMSE of 0.030, and a PCC of 0.968 (Fig. 2D). These results indicate that DECIPHER supports robust cellular composition inference across different molecular modalities within the same cellular system. In addition, DECIPHER is able to generalize to non-transcriptomic modalities. On the bone marrow metabolomic dataset, DECIPHER achieved a CCC of 0.784, an RMSE of 0.065, and a PCC of 0.808 on average (Fig. 2E), maintaining competitive performance and further supporting its applicability beyond transcriptomic data. On the breast proteomic dataset under the cross-physiological-state setting, DECIPHER achieved on average a CCC of 0.939, an RMSE of 0.032, and a PCC of 0.977 (Fig. 2F), showing slightly better performance than DECODE (CCC = 0.928, RMSE = 0.035, PCC = 0.971).

To further evaluate the robustness of DECIPHER, we stress-tested DECIPHER with four common types of perturbations by simulation, including random noise, systematic bias, feature missingness, and unknown cell types (Methods), on the PBMC CITE-seq RNA data and real PBMC bulk-cell transcriptomic datasets, namely Monaco, Microarray, and SDY67 (Methods; Fig. 2G, H). As a reference, we also conducted the same tests with DECODE. Changes in overall CCC were compared between DECIPHER and DECODE as the perturbation level increased. For the simulated bulk-cell data from the PBMC CITE-seq RNA dataset, both DECIPHER and DECODE maintained stable performance across the four perturbation types. DECIPHER retained higher CCC values under random noise, systematic bias, and unknown cell-type perturbations. Although performance gradually decreased with increasing feature missingness, the overall variation remained limited (Fig. 2G). A similar pattern was observed in the real PBMC bulk-cell datasets. Both methods showed relatively small fluctuations in overall CCC as the levels of random noise, systematic bias, feature missingness, and unknown cell-type perturbations increased (Fig. 2H). Feature missingness produced the most noticeable decrease in performance, but the overall changes remained moderate. Across both simulated and real-data settings, DECIPHER and DECODE therefore exhibited similar behavior after the four types of distinct perturbations. However, DECIPHER consistently achieved higher accuracy than DECODE across these conditions. Furthermore, using PBMC bulk-cell transcriptomic data from the Monaco, Microarray, and SDY67 cohorts, we evaluated DECIPHER under incomplete single-cell reference information (Methods; Fig. S1A-C). Cosine similarity analysis of the cell-type latent prototypes *Z*_ct_ revealed distinct similarity patterns among cell types, with relatively high similarity between CD8^+^ T and natural killer (NK) cells and partial similarity between CD4^+^ T and CD8^+^ T cells (Fig. S1B). Consistent with these similarity patterns, removal of the NK cell reference resulted in most of the missing signal being reassigned to CD8^+^ T cells, while removal of the CD4^+^ T cell reference similarly increased the predicted proportion of CD8^+^ T cells (Fig. S1A). Reference cell-type removal also affected deconvolution accuracy, with the magnitude of performance changes varying across cell types (Fig. S1C). These results suggest that the redistribution of predicted cell-type proportions following reference cell-type removal is associated with the similarity structure among *Z*_ct_ prototypes.

### Experimental validation of deconvolution performance in real bulk-cell mixtures

To further validate the deconvolution capability of DECIPHER under experimental conditions, we generated bulk-cell mixture datasets with known cell-type compositions (Fig. 3A; Methods). Ten cell lines were first profiled using single-cell RNA sequencing to construct the single-cell reference dataset (Fig. 3B-C). Subsequently, different cell lines were mixed according to predefined cell-type proportions and profiled using bulk RNA sequencing to generate experimentally measured bulk expression profiles (Fig. 3D). Each mixture design included nine experimental replicates, resulting in 45 bulk-cell samples for evaluating deconvolution performance across different mixture compositions and experimental replicates.

**Fig. 3:**
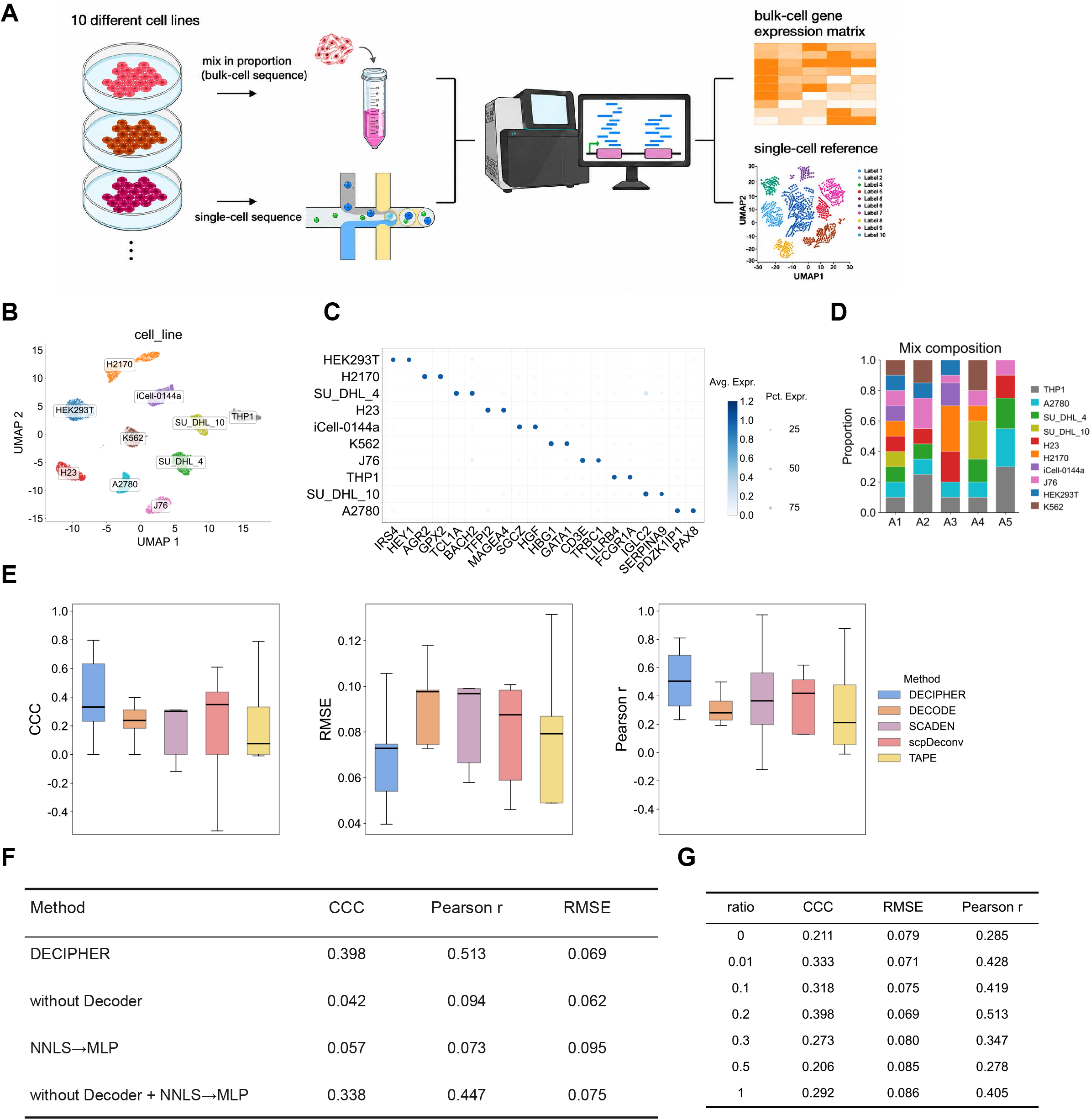
Evaluation of DECIPHER using experimentally generated bulk-cell mixtures and matched single-cell reference data. (A) Experimental design. Ten cell lines were profiled by single-cell RNA sequencing to construct the reference dataset. Different cell lines were mixed at predefined proportions and profiled by bulk-cell RNA sequencing to generate bulk-cell expression profiles for deconvolution. (B) UMAP (Uniform Manifold Approximation and Projection) visualization of the single-cell reference transcriptome profiles, color-coded by cell lines. (C) Dotplot showing expression of representative marker genes of the ten cell lines. Dot size indicates the proportion of cells expressing each marker, and color indicates the corresponding mean expression level. (D) Stacked bar plot showing the predefined compositions of the experimentally generated bulk-cell mixtures (A1–A5). (E) Comparison of DECIPHER with other deconvolution methods trained using simulated bulk-cell transcriptome profiles. Performance on the experimentally generated bulk-cell mixtures was evaluated using CCC, RMSE, and Pearson’s correlation coefficient (r). (F) Summary of ablation analysis results of DECIPHER on the experimentally generated bulk-cell RNA-seq dataset. The full DECIPHER model is compared with variants in which the decoder is removed, NNLS is replaced with a multilayer perceptron (MLP), or both modifications are applied. Performance was evaluated by CCC, Pearson’s correlation coefficient (r) and RMSE. (G) Summary of deconvolution performance of DECIPHER across different real-bulk Mixup ratios. The mixup ratio denotes the proportion of training inputs in the real bulk branch that are generated *ad hoc* by randomly mixing expression profiles from real bulk samples. Performance was evaluated by CCC, Pearson’s correlation coefficient (r) and RMSE.

Marker gene expression patterns supported the annotation of the ten cell lines (Fig. 3C), and the cell lines formed distinct clusters in the low-dimensional embedding (Fig. 3B). Principal component analysis (PCA) of the bulk-cell replicates further showed that samples from the same mixture design clustered closely together (Fig. S2A), indicating good consistency among experimentally generated bulk-cell profiles. On these experimental bulk-cell mixtures, DECIPHER recovered the predefined cell-type compositions with good agreement (Fig. S2B-C). We next compared DECIPHER with deep-learning-based deconvolution methods that learn a single, non-disentangled representation or directly map bulk expression profiles to cell-type proportions, rather than explicitly separating shared biological information from domain-associated variation. DECIPHER achieved the highest mean CCC on the experimental bulk-cell mixtures (approximately 0.40; Fig. 3E). In contrast, several competing approaches showed negative CCC values under certain mixture designs, indicating substantial deviations between predicted and true cell-type proportions. Given the limited number of mixture designs and the restricted variation in cell-type proportions, we additionally assessed overall concordance using the replicate-averaged proportions. Predicted and ground-truth proportions were pooled across the five mixture designs and ten cell types, yielding 50 paired values for the pooled CCC calculation. DECIPHER achieved the highest pooled CCC of approximately 0.78 among the compared methods (Fig. S2D). These results further highlight the advantage of DECIPHER’s disentangled representation design.

To investigate the contribution of individual model components, we performed ablation analyses (Fig. 3F; Methods). The full DECIPHER model achieved the highest CCC and correlation among the evaluated configurations. Removing the decoder or replacing NNLS with a multilayer perceptron (MLP) reduced concordance, whereas applying both modifications together partially recovered performance but remained inferior to the full model in terms of CCC and correlation. Although this configuration resembles the prediction strategy adopted by some deep learning-based deconvolution frameworks^13,14^, it remained inferior to the complete DECIPHER model. We further evaluated the effect of real data mix-up on model performance. During training, different proportions of mixed real bulk-cell expression profiles were introduced into the Real bulk-cell data branch (Methods). Different mix-up ratios resulted in distinct deconvolution performance, where the mix-up ratio represents the proportion of real-domain samples subjected to Mixup augmentation during training, with a ratio of 0.2 achieving the best performance (Fig. 3G). These results suggest that moderate augmentation of the Real bulk-cell data can improve DECIPHER performance on experimentally generated bulk data.

### Generalization across real and cross-modality datasets

To further evaluate the generalizability of DECIPHER across real biological datasets and cross-modality applications, we assessed its performance on real PBMC bulk-cell transcriptomic, spatial transcriptomic, and regulatory genomics datasets. These datasets represent diverse experimental sources, technological platforms, and molecular modalities, enabling evaluation of DECIPHER under complex biological conditions.

First, we evaluated DECIPHER on three real PBMC bulk-cell transcriptomic datasets, including Monaco^35^, Microarray^10^, and SDY67^36^ (Fig. 4A-C; Methods). DECIPHER maintained relatively stable performance across all three PBMC datasets and showed better overall performance than the other benchmark methods, with higher mean CCC and Pearson’s r and lower mean RMSE, indicating its ability to adapt to real bulk-cell transcriptomic profiles generated from different studies and experimental platforms. Using the same pooled evaluation strategy, DECIPHER also achieved the highest overall CCC in each of the three real PBMC datasets (Fig. S2E-G).

**Fig. 4:**
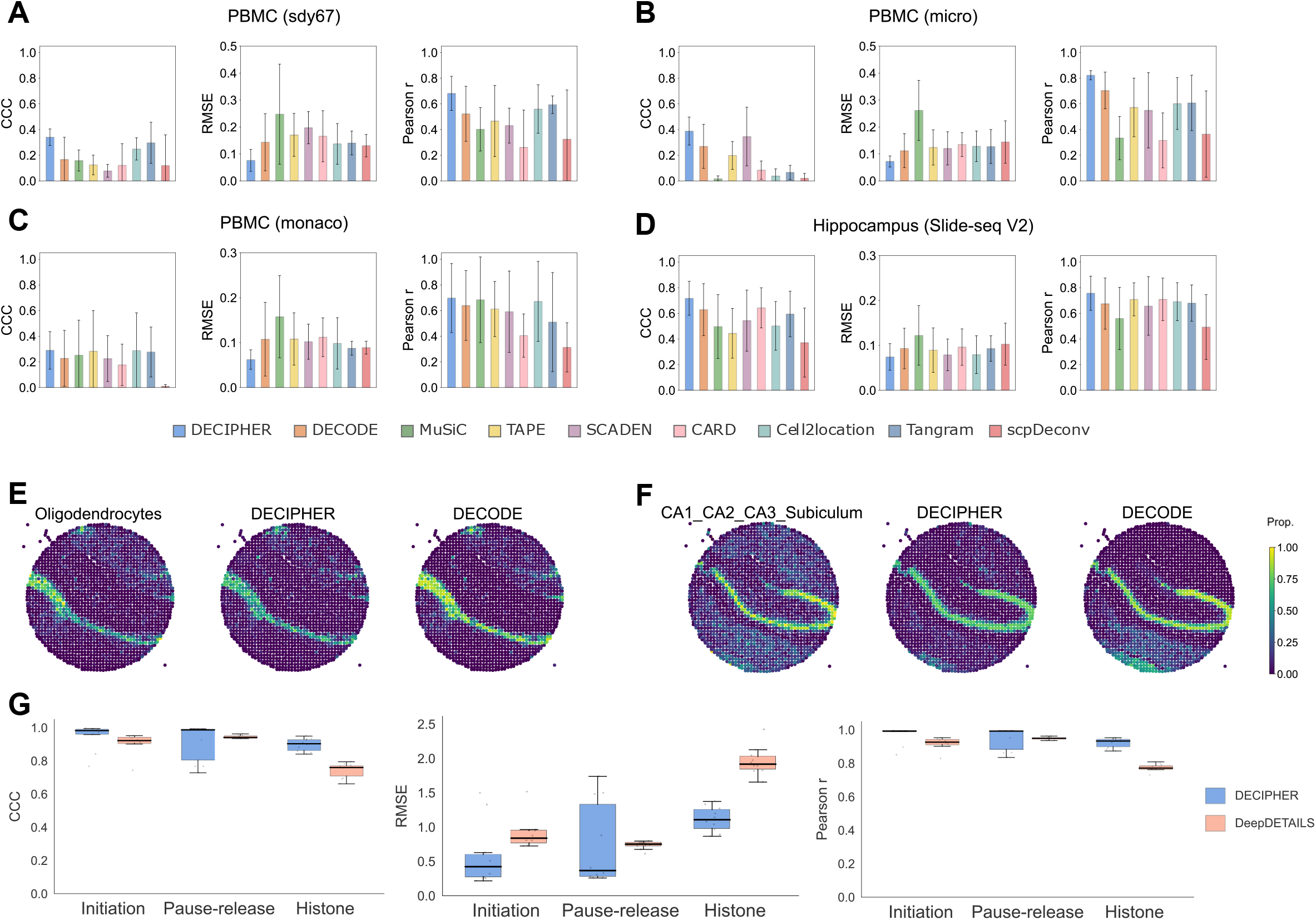
Performance of DECIPHER on real datasets and cross-modality deconvolution tasks. (A-D) Comparison of DECIPHER with other deconvolution methods on the SDY67 PBMC (A), PBMC microarray (B), PBMC Monaco (C) and mouse hippocampal Slide-seqV2 spatial transcriptomics (D) datasets, respectively. Deconvolution performance was evaluated using CCC, RMSE and Pearson’s correlation coefficient (r). (E-F) Spatial distributions of oligodendrocytes (E) and CA1/CA2/CA3/Subiculum neurons (F) in the mouse hippocampal Slide-seqV2 spatial transcriptomics dataset. Ground truth cell-type proportions (left) and corresponding predictions from DECIPHER (middle) and DECODE (right) are shown. The color scale indicates the cell-type proportion at each spatial location, with higher values corresponding to greater relative abundance of the indicated cell type. (G) Comparison of DECIPHER and DeepDETAILS for cross-modality deconvolution using single-cell ATAC-seq data as the reference. Deconvolution results are shown for transcription initiation, pause-release, and histone modification datasets and evaluated using CCC, RMSE, and Pearson’s correlation coefficient (r).

We next evaluated DECIPHER on spatial transcriptomic data using the mouse hippocampal Slide-seqV2 dataset^37^ (Methods). DECIPHER inferred spatial cell-type compositions with a mean CCC of 0.717, an RMSE of 0.074, and a PCC of 0.757 across spatial grids (Fig. 4D). As representative examples, oligodendrocytes and CA1/CA2/CA3/Subiculum neurons were selected to visualize the inferred spatial distributions, which showed good agreement with the reference proportions (Fig. 4E-F). Notably, the local spatial distributions inferred by DECIPHER appeared relatively stable and clean, suggesting that DECIPHER may have a certain degree of denoising capability in spatial transcriptomic data.

Finally, we compared DECIPHER with DeepDETAILS^38^ for cross-modality regulatory genomics deconvolution (Methods; Fig. 4G). Using single-cell profiles generated by the assay for transposase-accessible chromatin using sequencing (ATAC-seq) as references, DECIPHER was evaluated for the deconvolution of three regulatory processes: transcription initiation, pause release, and histone modification. Performance was evaluated on the recovered cell-type-specific regulatory signals after the second-stage signal-recovery procedure. DECIPHER achieved high performance across all three regulatory stages. For transcription initiation, DECIPHER achieved a mean CCC of 0.947, an RMSE of 0.586, and a PCC of 0.969. The corresponding values were 0.913, 0.743, and 0.947 for pause release, and 0.897, 1.114, and 0.923 for histone modification. In comparison, DeepDETAILS achieved mean CCC values of 0.909, 0.945, and 0.744 for the corresponding stages, respectively, with DECIPHER showing higher concordance particularly for histone modification prediction. On the same cross-modality tasks, DECIPHER required an average runtime of approximately 0.13 h, including signal recovery, compared with 14.9 h for DeepDETAILS (Fig. S2H; Methods), indicating substantially higher computational efficiency while maintaining competitive predictive performance.

### DECIPHER representations support immune-age prediction across PBMC datasets

To further evaluate whether the latent representations learned by DECIPHER contain biologically meaningful information, we collected multiple PBMC bulk-cell RNA sequencing (RNA-seq) datasets with age information and deconvolved them using DECIPHER. The resulting *Z*_c_ was used for immune age prediction, as PBMCs consist primarily of immune cells and their cellular composition varies across the human lifespan^21^. *Z*_s_ was included as a comparison to assess age-associated information retained in two latent spaces. We first visualized the *Z*_c_ and *Z*_s_ representations learned by DECIPHER using uniform manifold approximation and projection (UMAP)^39^, with samples color-coded according to batch (i.e., data source) information (Fig. 5A). The *Z*_s_ latent space showed more pronounced batch-associated structure (Fig. 5A right), whereas such structure was less apparent in *Z*_c_ (Fig. 5A left). The latent representation learned by DECODE also showed more evident batch-associated structure (Fig. 5B).

**Fig. 5:**
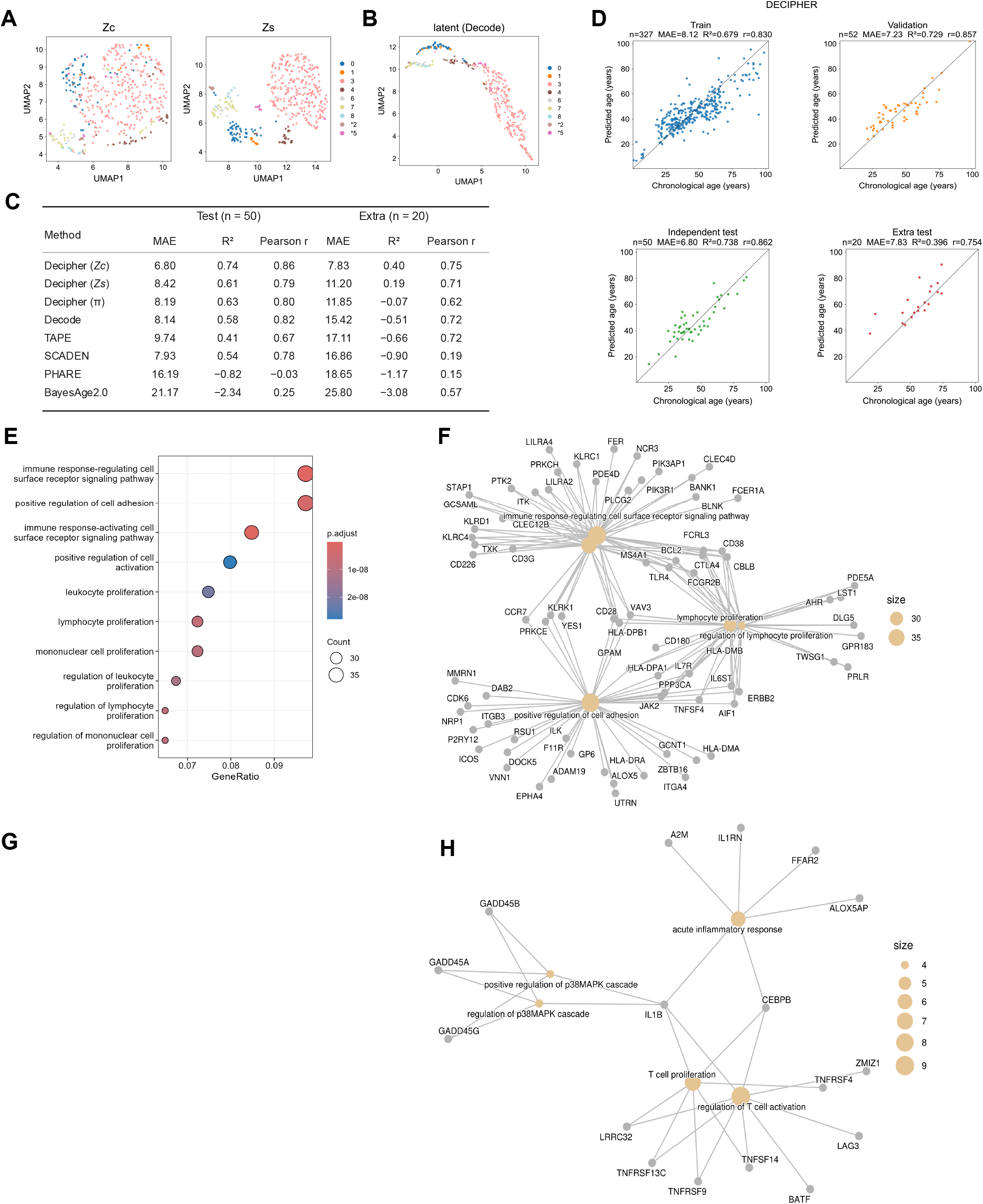
Chronological age prediction from PBMC bulk-cell RNA-seq profiles using DECIPHER latent representations. (A) UMAP projections of the *Z*_c_ (left) and *Z*_s_ (right) representations learned by DECIPHER, colored by batch (data source). Asterisks denote samples from the test batch. (B) UMAP projection of the latent representation learned by DECODE, colored by batch (data source). (C) Comparison of age-prediction performance using DECIPHER-derived latent representations (*Z*_c_ and *Z*_s_), estimated cell-type proportions (π), other representation-learning models, and established age-prediction methods. Performance is evaluated using mean absolute error (MAE), coefficient of determination (R²), and Pearson’s correlation coefficient (r). Lower MAE indicates smaller prediction errors, whereas higher R² and Pearson’s correlation (r) indicate better predictive performance. (D) Age-prediction performance based on the DECIPHER domain-constant representation *Z*_c_. Scatterplots show predicted versus chronological age across the training, validation, independent test, and additional test datasets. The dashed lines represent the line of identity. (E) Gene Ontology (GO) Biological Process enrichment analysis of genes associated with the age-associated latent feature *Z*_c18_ (the 18^th^ dimension in *Z*_c_), which exhibits the strongest negative Pearson correlation with age among the latent features in *Z_c_*. Genes with expression levels positively correlated with *Z_c_*18 were selected for GO Biological Process enrichment analysis. In the dot plot, the x-axis indicates GeneRatio, dot size represents the number of genes associated with each GO term, and color denotes the adjusted P value. (F) Gene–GO term network showing the relationships between enriched GO terms and their associated genes in (E). (G) Same as E, but for genes negatively correlated with *Z*_c18_. (H) Gene–GO term network showing the relationships between enriched GO terms and their associated genes in (G).

Next, we used the *Z*_c_ representation learned by DECIPHER for immune-age prediction and compared its performance with *Z*_s_, the estimated cell-type proportions (π), existing immune-age prediction methods, and other deconvolution-based representations (Fig. 5C). Prediction results for *Z*_c_, *Z*_s_, and π are shown in Fig. 5D and Fig. S3A,B, respectively. DECIPHER-*Z*_c_ achieved MAEs of 6.80 and 7.83 in the independent and additional test datasets, respectively, compared with 8.42 and 11.20 for *Z*_s_ and 8.19 and 11.85 for π. Overall, *Z*_c_ showed the best predictive performance and generalization across the evaluated datasets. Together with the UMAP patterns, these results suggest that *Z*_c_ retains more transferable age-associated information with reduced batch-associated structure, whereas *Z*_s_ retains more dataset-specific variation, further highlighting the advantage of DECIPHER’s disentangled representation design in separating transferable biological signals from domain-specific variation.

### Age-associated latent features capture biologically informative immune signals

To further characterize the age-associated information encoded in *Z*_c_, we first performed correlation analysis between the 32 *Z*_c_ latent features and age to identify age-associated features (Fig. S3C). We then further examined the gene expression patterns associated with the six *Z*_c_ features that most strongly correlated with age (Fig. S3D). The correlation heatmap revealed structured patterns between latent features and their associated genes, indicating that different dimensions of the *Z*_c_ latent space correspond to distinct gene expression changes. To further examine the information captured by these age-associated features, we compared gene sets associated with the same six *Z*_c_ dimensions, their composition-reconstructed representation *Z_c_*__recon_ = *wZ_ct_*, and the reconstruction residual *Z’*_c_ = *Z̄*_c_ − *Z_c_*__recon_ (Methods; Fig. S3E). Among these genes, 609 were shared by *Z*_c_ and *Z*^’^*_c_* but not *Z* and were mainly enriched in leukocyte activation, migration, chemotaxis, and related immune processes (Fig. S3F). In contrast, the 104 genes shared by *Z*_c_, *Z_c_*__recon_, and *Z*^′^ were enriched in platelet activation, coagulation, and cell-adhesion-related processes (Fig. S3G). These results indicate that the age-associated *Z_c_* features retain biological information that is not fully captured by the composition-reconstructed component.

We then selected *Z*_c18_, the latent feature most strongly associated with age, and performed Gene Ontology (GO) Biological Process enrichment analysis^40^ of its associated genes (Fig. 5E-H). Genes positively correlated with this latent feature were mainly enriched in immune response-regulating cell surface receptor signaling, positive regulation of cell adhesion, immune response-activating cell surface receptor signaling, and immune cell proliferation-related processes. In contrast, negatively correlated genes were mainly enriched in regulation of T cell activation, and acute inflammatory response. This gene set included LAG3 and BATF (Fig. 5H), which have been implicated in the regulation of T cell exhaustion^41,42^. These findings link age-associated *Z*_c18_ to immune activation, proliferation, and inflammatory regulation, with potential connections to exhaustion-associated transcriptional programs, supporting the biological interpretability of the DECIPHER *Z*_c_ representation.

Together, these results show that *Z*_c_ not only supports cell-type proportion estimation but also retains biological information associated with immune age and shows good generalization for age prediction across PBMC datasets. Its age-associated latent features were further linked to multiple immune regulatory processes, supporting the use of the DECIPHER *Z*_c_ latent representation for downstream phenotype analyses beyond deconvolution.

#### Prognostic stratification using latent representations of DECIPHER in lung adenocarcinoma

To further evaluate the utility of DECIPHER latent representations for disease-related phenotype modeling, we extracted *Z*_c_ trained using The Cancer Genome Atlas lung adenocarcinoma (TCGA-LUAD)^43^ RNA-seq dataset to construct a prognostic risk score and evaluate its association with patient survival outcomes (Fig. 6; Methods).

**Fig. 6:**
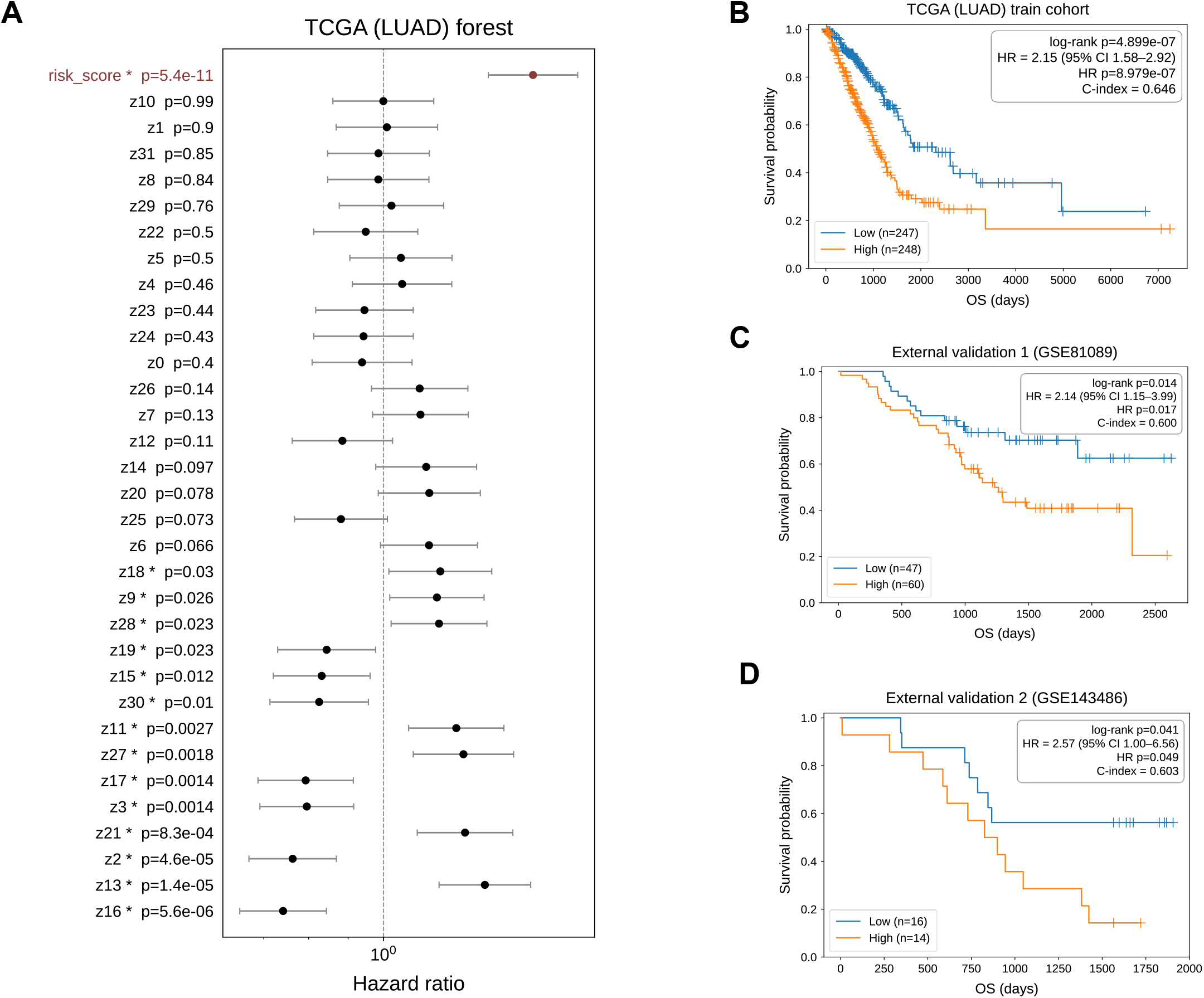
Prognostic evaluation using DECIPHER latent representations in lung adenocarcinoma. (A) Forest plot summarizing Cox proportional hazards estimates for the DECIPHER-derived risk score and individual latent features (*Z*_c_) in the TCGA-LUAD (lung adenocarcinoma) cohort. Points indicate hazard ratios (HRs), horizontal lines indicate 95% confidence intervals (CIs), and the dashed vertical line denotes HR = 1. HR > 1 indicates an association with an increased hazard of death, whereas HR < 1 indicates an association with a decreased hazard. A 95% CI that includes 1 indicates that the estimated association is not statistically significant at the corresponding confidence level. P values are shown for each variable. (B–D) Kaplan–Meier overall survival curves for the high- and low-risk groups defined using DECIPHER-derived risk scores in the TCGA-LUAD training cohort (B) and the external validation cohorts GSE81089 (C) and GSE143486 (D). Follow-up time in days is plotted against overall survival probability. Tick marks indicate censored observations, and n denotes the number of patients in each risk group. The log-rank P value assesses the difference in survival distributions between the two groups. HR denotes the hazard of death in the high-risk group relative to that in the low-risk group and is reported with its 95% CI. The concordance index (C-index) quantifies the ability of the model to discriminate between patients with different survival risks; values closer to 1 indicate better discrimination, whereas a value of 0.5 indicates performance comparable to random prediction.

We first performed Cox proportional hazards regression^44^ using the *Z*_c_ latent features learnt from the TCGA-LUAD samples and constructed a risk score (Fig. 6A). Multiple *Z*_c_ features were significantly associated with patient mortality risk, and the *Z*_c_-based risk score was significantly associated with overall survival. Patients were subsequently stratified according to the risk score, and Kaplan-Meier survival analysis^45^ was performed to evaluate differences in overall survival between risk groups (Fig. 6B). We next applied the TCGA-LUAD-derived risk model, without refitting, to two external validation RNA-seq cohorts, GSE81089^46^ and GSE143486^47^ (Fig. 6C-D). The risk score remained prognostic in both cohorts (GSE81089: HR = 2.14, 95% CI = 1.15-3.99; C-index = 0.600; GSE143486: HR = 2.57, 95% CI = 1.00-6.56; C-index = 0.603). In contrast, risk scores derived from *Z*_s_ showed limited prognostic stratification ability across TCGA-LUAD and the external validation cohorts (Fig. S4A-C). Risk scores based on the estimated cell-type proportions (π) also showed weaker prognostic stratification, particularly in the external validation cohorts (Fig. S4D-F).

To further characterize the biological information encoded in the prognostic *Z*_c_ representation, we selected the six *Z*_c_ features most significantly associated with survival outcomes and examined their relationships with gene expression profiles (Fig. S4G). We then selected *Z*_c16_, the latent feature with the largest absolute Cox regression coefficient among the standardized features (Fig. 6A), and performed GO enrichment analysis of its associated genes (Fig. S4H-I). The enriched biological processes were mainly related to adaptive immune responses, B-cell receptor signaling, and regulation of immune responses. To further examine whether this prognostic information could be fully explained by estimated cellular composition, we compared gene sets associated with the same six prognosis-associated *Z_c_* dimensions, their composition-reconstructed representation *Z_c_*__*recon*_, and the reconstruction residual *Z^’^_c_* (Fig. S4J). Among these genes, 240 were shared by *Z_c_* and *Z^’^_c_* but not *Zc_recon* and were mainly enriched in B-cell receptor signaling, immunoglobulin-mediated immune responses, and related humoral immune processes (Fig. S4K). In contrast, the 10 genes shared by *Z_c_*, *Z_c_*__*recon*_, and *Z*^′^ were enriched mainly in T-cell activation, differentiation, and related immune-regulatory processes (Fig. S4L). These results indicate that the prognostic *Z_c_* representation retains biologically informative variation that is not fully captured by the composition-reconstructed component.

Together, these results show that *Z*_c_ captures prognostic information in LUAD, supporting both risk stratification in external cohorts and the analysis of biological factors associated with survival outcomes. These findings further support its use beyond cell-type deconvolution as a biologically informative, domain-constant representation for downstream phenotype analyses.

## Discussion

Our results show that DECIPHER achieves robust cell-type deconvolution across simulated datasets, experimentally generated bulk-cell mixtures, and real-world datasets while providing a unified modeling framework applicable across multiple molecular modalities. Extensive benchmarking demonstrated that DECIPHER outperformed existing deconvolution methods in nearly all tested scenarios. More importantly, the cell-related representation *Z*_c_ learned during deconvolution captured biologically informative variation beyond cell-proportion estimation, as supported by its stronger performance than the estimated cell-type proportions *π*in immune-age prediction and survival analysis. These results establish that the latent representations learned by DECIPHER support not only cell-type proportion estimation but also broader representation learning from bulk molecular data.

An important feature of DECIPHER is the integration of deep representation learning with interpretable proportion estimation through an explicit compositional constraint. Neural-network regressors optimized directly for cell proportions may learn features that are sufficient for prediction but do not necessarily preserve broader molecular characteristics of bulk-cell samples^13,14,20^. DECIPHER instead integrates differentiable NNLS optimization^48^ into the latent space learned by an autoencoder, modeling each sample’s cell-composition representation as a non-negative combination of cell-type reference prototypes. This design maintains an interpretable relationship between latent features and cellular composition while avoiding a fully black box regression mapping. Beyond this compositional constraint, DECIPHER further benefits from its disentangled representation design. Its superior performance on experimentally generated real-bulk mixtures, together with the stronger performance of *Z_c_* than *Z_s_* and the estimated cell-type proportions *π* in immune-age prediction and survival analysis, supports the advantage of this design in preserving transferable biological information while separating domain-specific variation.

Generalization from simulated data to real bulk-cell data also presents a practical challenge because many simulated mixtures can be generated, whereas the availability of real target samples is often relatively limited. Consequently, even with cross-domain learning, the distribution on the real-data side may be defined by only a small number of discrete samples, increasing the risk that the model becomes overly dependent on the limited target-domain observations^14,16,25^. To increase sample diversity on the real-data side, DECIPHER applies Mixup augmentation to real bulk-cell profiles during training. Rather than simply increasing the number of training samples, this strategy constructs intermediate target-domain expression profiles between observed real samples, thereby increasing the coverage density of the real-bulk expression space and extending the target domain from relatively discrete observations toward a more continuous expression space. This augmentation can reduce overfitting to individual real samples and provide a smoother and more diverse target-domain representation for alignment between simulated and real data. Comparisons across different Mixup ratios further demonstrated that moderate expansion of target-domain coverage improved model performance, whereas stronger mixing did not continuously provide additional benefit, indicating that an appropriate balance is required between increasing target-domain diversity and preserving the structure of the original real samples.

Several limitations remain in our study. First, although DECIPHER can be applied across multiple molecular modalities, the current implementation remains task-specific and requires separate model training for individual reference-target settings rather than allowing a single trained model to be directly transferred across tissues, platforms, or molecular modalities. When substantial batch effects, technical discrepancies, or modality differences exist between reference and target datasets, dataset-specific training or adaptation may therefore still be required^27^. Second, model performance remains dependent on the quality and cell-type coverage of the single-cell reference^3^, particularly when target samples contain rare cell populations or substantially altered cellular states. A promising future direction is to develop more generalizable pre-trained frameworks^23,24^ using larger collections of cross-tissue, cross-platform, and cross-omics datasets, allowing models to learn more transferable biological representations while reducing the need for dataset-specific retraining. However, pretraining alone does not guarantee generalization across biological and technical conditions, and task- or dataset-specific fine-tuning may still be required. Moreover, evaluations of foundational models have revealed limitations in batch integration relative to established methods fitted to individual datasets^49^. Dataset-specific frameworks such as DECIPHER therefore remain valuable for explicitly modeling reference-target differences while jointly estimating cellular composition and learning biologically informative representations for downstream analysis.

In summary, DECIPHER combines nonlinear representation learning, explicit modeling of shared and domain-associated variation, and differentiable NNLS optimization within an end-to-end framework for cell-type deconvolution. DECIPHER demonstrates competitive deconvolution performance across molecular modalities, with *Z*_c_ supporting downstream phenotype modeling and biological interpretation. DECIPHER thus connects cellular composition analysis with biologically informative sample representation learning, broadening the utility of existing bulk omics resources for biological and clinical research.

## Methods

### DECIPHER model

#### Overall model framework

DECIPHER jointly trains on simulated mixtures with known cell-type proportions and unlabeled target samples. Both sample types share the same model parameters. Simulated samples provide direct cell-proportion supervision, whereas simulated and target samples jointly contribute to reconstruction and domain-related objectives.

For an input profile *x_i_*, a shared encoder *E_θ_* first produces an intermediate representation ℎ*_i_*,

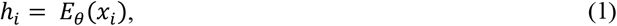

which is subsequently mapped by separate projection heads into a domain-constant representation *Z*_c,*i*_ and a domain-specific representation *Z*_s,*i*_,

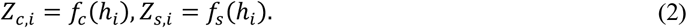

The complete *Z*_c_ and *Z*_s_ representations are concatenated and passed to a decoder to reconstruct the input molecular profile.

### Cell-type reference prototypes and differentiable NNLS deconvolution

Cell-type reference representations were derived from the single-cell reference data to characterize the latent representation of each reference cell type. Each reference prototype was L2-normalized and mapped through a learnable projection network into the same latent space as the sample-level domain-constant representation **Z***_c_*_,*i*_. The projected cell-type prototypes were further L2-normalized to obtain **Z***_ct_* ∈ ℝ*^C^*^×*d*^, whose rows correspond to the *C* reference cell types. To distinguish the original sample representation from the representation used for NNLS fitting, the L2-normalized sample representation was defined as

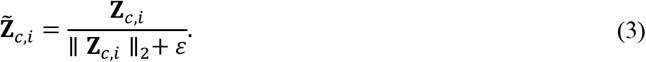

Throughout this study, *π* denotes a generic cell-type proportion vector. When known and model-estimated proportions need to be distinguished explicitly, *π*^true^ and *π*. denote the known and model-estimated proportions, respectively.

For each sample, 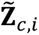 was decomposed against the projected cell-type prototypes using a differentiable non-negative least-squares (NNLS) objective^50^:

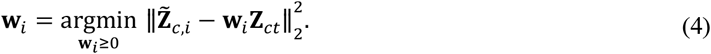

The NNLS problem was implemented using a differentiable mirror-descent optimization procedure^50^. The resulting non-negative coefficients were normalized to obtain the estimated cell-type fractions:

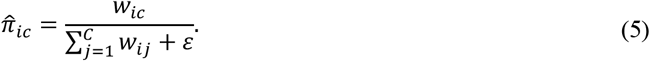

where C denotes the number of reference cell types and ε is a small constant for numerical stability.

### Loss functions

DECIPHER was optimized using six consistently defined objectives: the cell-proportion supervision loss (ℒ_prop_), input reconstruction loss (ℒ_rec_), latent reconstruction loss (ℒ_latrec_), domain classification loss (ℒ_dom_), cross-domain alignment loss (ℒ_align_), and proportion-aware contrastive loss (ℒ_contrast_). The total loss was thus

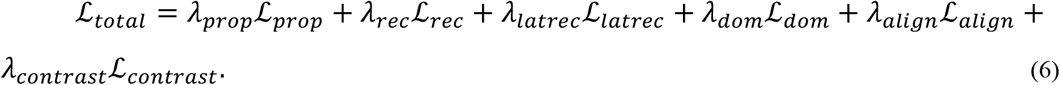

For simulated samples with known cell-type proportions, the cell-proportion supervision loss combined mean squared error (MSE) and soft-label cross-entropy (CE):

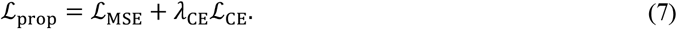

where

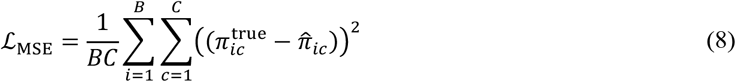

and

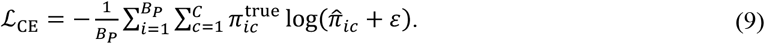

The CE contribution was dynamically scaled from exponential moving averages (EMA) of the two components. For x ∈ {MSE, CE},

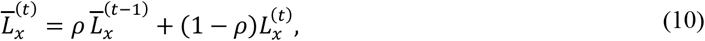

and 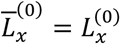. The scaling weight was

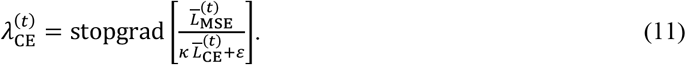

Here, the EMA decay coefficient was set to *ρ* = 0.99, and *κ* denotes the MSE-to-CE balancing ratio. A value of *κ* = 20 was used for the standard deconvolution benchmarks, whereas *κ* = 4 was used in the immune-age analysis.

The input reconstruction loss was calculated for both simulated (*P*) and target (*R*) samples:

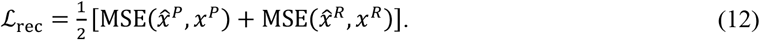

For the gated latent *Z*_c_ reconstruction loss, the per-sample prototype reconstruction error and coefficient entropy were defined as

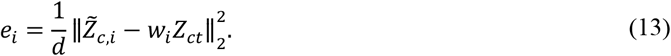

Here, *H_i_* measures the diversity of the predicted cell-type composition rather than predictive uncertainty.

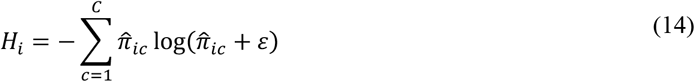

The detached gate was defined as

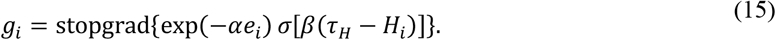

The gated latent reconstruction loss was then

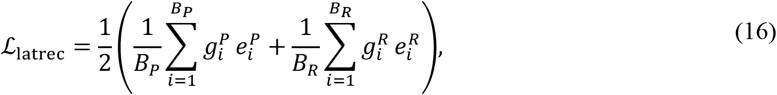

with *α* = 5, *β* = 10, and *τ*_H_ = 2.

Domain information was supervised through the domain classification loss on the domain-specific representation *Z*_s_:

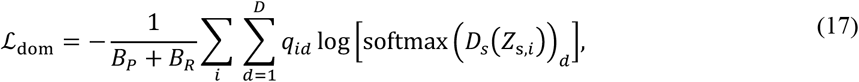

where *q_id_* denotes the domain label and *D_s_* denotes the domain classifier.

Cross-domain alignment was imposed in the cell-composition latent space using maximum mean discrepancy (MMD)^51^:

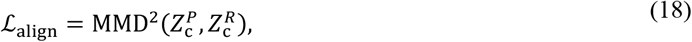

For the proportion-aware contrastive loss, cosine similarity between the true proportion vectors of simulated samples *i* and *j* was defined as

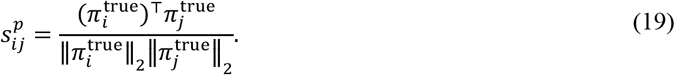

The positive-sample set for anchor i was defined as

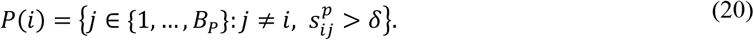

with δ = 0.9. After L2 normalization of *Z*_c_, latent similarity was defined as

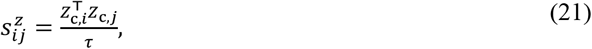

where *τ* = 0.2 is the temperature parameter. Let

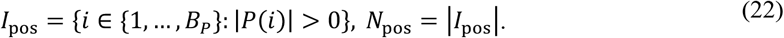

For compact notation, define

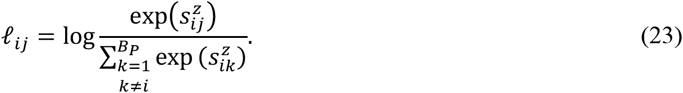

The proportion-aware contrastive loss, computed only over simulated samples, was

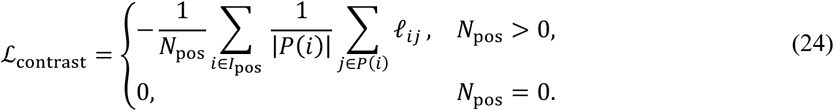

The default loss weights were set to (100, 0.1, 0.1, 0.2, 0.3, 0.1), corresponding to (*λ_prop_, λ_rec_, λ_latrec_, λ_dom_, λ_align_, λ_contrast_*), with κ = 20. For Slide-seqV2 hippocampal analysis, λalign was set to 0.01. For the immune-age analysis, *κ* = 4was used.

### Model training

Models were optimized using AdamW^52^ with a learning rate of 1 × 10^−4^, weight decay of 1 × 10^−3^, batch size of 128, and a maximum of 300 epochs. Early stopping with a patience of 10 epochs was applied according to validation loss. The implementation used Python 3.8 and PyTorch 2.0.0^53^ with CUDA 11.8 on an NVIDIA RTX 4090 GPU.

### Comparison with DECODE

Compared with DECODE^15^, DECIPHER differs in four aspects: (i) it jointly optimizes representation learning and deconvolution within an end-to-end framework, whereas DECODE performs domain adaptation and subsequent feature denoising in sequential training stages; (ii) DECODE separates purified features from noise features through attention-based denoising and contrastive learning, whereas DECIPHER explicitly learns a domain-constant representation (*Z*_c_) and a domain-specific representation (*Z*_s_), guided by cross-domain alignment and domain-label prediction, respectively; (iii) decoder-based input reconstruction to encourage information retention beyond cell-proportion supervision; and (iv) differentiable NNLS estimation using reference-derived cell-type prototypes, providing an explicit compositional constraint instead of direct neural network regression.

### Benchmarking and model evaluation

#### Comparative methods and evaluation metrics

DECIPHER was compared with DECODE^15^, MuSiC^8^, TAPE^14^, SCADEN^13^, CARD^19^, Cell2location^17^, Tangram^18^, and scpDeconv^16^. When permitted by the input requirements of each method, the same reference/test partitions and matched feature spaces were used.

Deconvolution performance was evaluated using Lin’s concordance correlation coefficient (CCC)^28^, Pearson’s correlation (*r*), and root mean squared error (RMSE). For a given cell type, *π*^true^ and *π*. denote the observed and model-predicted proportion vectors, respectively. CCC was calculated as

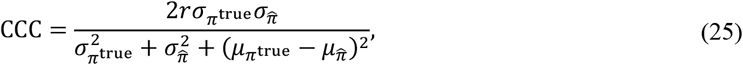

where μ and σ denote the corresponding means and standard deviations of the true and predicted proportion vectors. Pearson’s correlation was calculated as

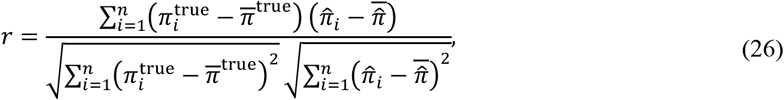

and RMSE was calculated as

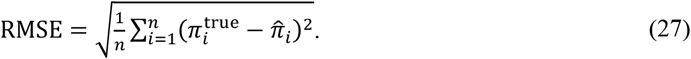

Unless otherwise specified, metrics were calculated for each cell type across samples and then averaged across cell types. For the experimental bulk-cell mixtures, predictions were first averaged across the nine replicates within each of the five mixture designs. Metrics in Fig. 3 were calculated across cell types within each mixture and then averaged across mixtures, whereas the pooled CCC in Fig. S2 was calculated after flattening the replicate-averaged proportions across all five mixtures and ten cell types (n = 50).

### Robustness and incomplete-reference analyses

Robustness was evaluated using CITE-seq RNA and the real PBMC bulk-cell datasets. Four perturbation types were examined at intensities of 0%, 5%, 10%, 15%, and 20%: random expression noise, systematic expression bias, feature missingness, and unmodeled cellular-state contamination. Random expression noise was introduced by multiplying selected features by independent factors sampled from *Uniform*(0.9,1.1) . Systematic bias was introduced by multiplying selected features by either 0.8 or 1.2. Feature missingness was simulated by setting selected features to zero. Unmodeled cellular-state contamination was generated by combining the mean profiles of two randomly selected reference cell types at a 1:1 ratio and introducing the resulting profile at the specified intensity.

Reference incompleteness was further evaluated using leave-one-cell-type-out analysis. One cell type was removed from the reference at a time, deconvolution was repeated, and changes in predicted proportions and performance were recorded. Cosine similarity between the omitted cell-type prototype and the retained *Z*_ct_ prototypes was calculated to characterize redistribution toward similar retained cell types.

### Real-bulk Mixup augmentation

All 45 experimentally generated real-bulk samples were retained in the real domain. Before Mixup, raw bulk-cell counts were processed by CPM (counts per million) scaling, log1p transformation, and sample-wise maximum normalization to obtain normalized expression vectors. During training, Mixup augmentation was applied to real-domain samples with a probability of 20%. When applied, several real-bulk expression profiles were randomly selected, and a set of non-negative weights summing to one was sampled from a symmetric Dirichlet distribution with concentration parameter *α* = 0.5. The normalized expression vectors were then linearly combined using these weights to generate intermediate expression states between the observed real-bulk samples. This augmentation was used to increase the diversity of the real-domain training distribution and limit overfitting to the finite set of real samples. Mixup was disabled during validation and evaluation, which were performed using the original processed real-bulk profiles.

### Experimental generation and sequencing of bulk-cell mixtures

The experimental bulk-cell mixture benchmark was generated using ten human cell lines: THP1, A2780, SU-DHL-4, SU-DHL-10, NCI-H23, NCI-H2170, iCell-0144a, J76, HEK293T, and K562. Detailed source information for the six cell lines maintained in our laboratory (SU-DHL-4, SU-DHL-10, NCI-H23, NCI-H2170, iCell-0144a, and K562) is provided in Supplementary Table 2. THP1, A2780, J76, and HEK293T were kindly provided by the laboratory of Guobing Chen. Cells were mixed according to five predefined composition designs (A1-A5; Fig. 3D). Each predefined mixture was divided into nine aliquots, and each aliquot underwent independent RNA extraction and bulk-cell RNA sequencing, resulting in 45 experimental bulk-cell RNA-seq samples in total. The predefined mixing proportions were used as ground truth for evaluating deconvolution performance. Total RNA was extracted using TRIzol reagent according to the manufacturer’s protocol. RNA quantity and purity were assessed using NanoDrop 2000, and RNA integrity was evaluated using an Agilent 2100 Bioanalyzer. Bulk-cell RNA-seq libraries were constructed using the VAHTS Stranded mRNA-seq Library Prep Kit for Illumina V2 (Vazyme) and sequenced on the DNBSEQ-T7 platform.

Raw sequencing reads were evaluated using FastQC^54^, followed by adapter and low-quality read trimming using Trim Galore^55^. Filtered reads were aligned to the human reference genome (hg38) using HISAT2^56^, and gene-level counts were quantified using featureCounts^57^. The resulting count matrices were converted to CPM and log1p transformed before downstream analysis.

A matched single-cell RNA-seq reference was generated from the same ten cell lines using the Chromium GEM-X Single Cell 3′ Reagent Kit v4 (10x Genomics)^58^ following the manufacturer’s protocol. The processed single-cell reference was used to generate simulated bulk-cell profiles for supervised DECIPHER training.

### Downstream representation analyses

#### Immune-age prediction and interpretation

We collected 449 healthy PBMC bulk RNA expression samples for the immune-age analysis. Two complete study batches (batch 2 and batch 5; *n* = 20 in total) were held out as an additional test set and were not used for training, validation, or model selection of the age-prediction model, but were used only for external evaluation. These held-out batches were included as unlabeled target-domain samples during DECIPHER representation learning but were not used for training, validation, or model selection of the downstream age-prediction model. The remaining 429 samples were stratified into 10-year age bins according to chronological age and randomly assigned within each age bin, resulting in 327 training samples, 52 validation samples, and 50 independent test samples. This stratified splitting procedure was used to maintain approximately balanced and comparable age distributions across the three partitions.

The *Z_c_* features were used as input to a multilayer perceptron regressor with hidden layers of 64 and 32 units, ReLU activation, and dropout of 0.1. The age-prediction model was optimized using AdamW with a learning rate of 1 × 10^−3^, weight decay of 1 × 10^−4^, batch size of 32, a maximum of 400 epochs, and early stopping with a patience of 40 epochs according to validation MAE. An L1 objective was used for training. For comparison, the same age-prediction workflow was applied separately to *Z*_s_ and the estimated cell-type proportion vector π. Age-prediction performance was evaluated using mean absolute error (MAE), coefficient of determination (*R*^2^), and Pearson’s correlation (*r*) between predicted and chronological age.

For biological interpretation of *Z_c_*, Pearson’s correlation (*r*) was first calculated between each *Z_c_* dimension and chronological age. *Z_c_* dimensions satisfying ∣ *r* ∣> 0.20 were defined as age-associated dimensions. Candidate dimensions were ranked according to the absolute value of age correlation, and the top six dimensions were selected for downstream gene-association analysis. For each selected age-associated *Z_c_*dimension, Pearson’s correlation (*r*) was calculated between the latent dimension and gene expression levels. Genes were ranked according to the absolute correlation coefficient, and genes with the strongest associations were selected for correlation visualization. Positively and negatively correlated gene sets were separately analyzed for Gene Ontology (GO) Biological Process enrichment using the R package clusterProfiler^59^. The 1,295 genes included in the correlation analysis were used as the background gene set, and enrichment significance was assessed using the Benjamini-Hochberg method for multiple-testing correction.

### Lung-cancer representation and prognostic analysis

TCGA-LUAD and TCGA-LUSC bulk transcriptomic samples were jointly included as target-domain samples during lung-cancer representation learning. Prognostic modeling was subsequently performed in TCGA-LUAD, with the lung adenocarcinoma cohorts GSE81089 and GSE143486 used for external validation. GSE81089 and GSE143486 were included as unlabeled target-domain samples during DECIPHER representation learning, whereas the Cox model and risk threshold were estimated exclusively from TCGA-LUAD and applied to these cohorts without refitting. A ridge Cox proportional-hazards model^44^ was then fitted using all standardized *Z*_c_ features. For sample *i*, the risk score was defined as

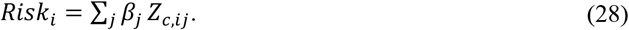

where *β*_j_ denotes the Cox regression coefficient estimated in TCGA-LUAD for the *j*th *Z*_c_ feature. No *Z*_c_ feature was excluded from risk-score construction solely on the basis of its individual statistical significance.

TCGA-LUAD patients were divided into high- and low-risk groups using the median TCGA-LUAD risk score as the cutoff. For external validation, risk scores for GSE81089 and GSE143486 were calculated using the fixed Cox coefficients learned in TCGA-LUAD. The TCGA-derived median cutoff was applied unchanged in both external cohorts without refitting the Cox model or re-estimating the threshold. Survival differences were evaluated by Kaplan-Meier analysis^45,46^ and the log-rank test, and hazard ratios with 95% confidence intervals and concordance indices were reported. For comparison, the same prognostic workflow was applied separately to *Z*_s_ and π.

For biological interpretation of the prognostic representation, Pearson’s correlation (*r*) was calculated between prognostic *Z_c_* features and gene expression in TCGA-LUAD. Genes associated with prognostic *Z_c_* features were selected using an absolute correlation threshold of ∣*r* ∣> 0.4 and categorized into positively and negatively correlated gene sets according to correlation direction. Because some *Z_c_* features (e.g., *Z_c_*_16_ shown in Fig. S4) were associated with only a limited number of negatively correlated genes, Gene Ontology Biological Process enrichment analysis was primarily performed on the positively correlated gene sets.

### Decomposition of the ***Z_c_*** representation

To distinguish the component of *Z_c_* explained by the estimated cell-type composition from the remaining representation, we reconstructed the cell-composition-associated component using the estimated cell-type proportion vector and the corresponding cell-type latent prototypes:

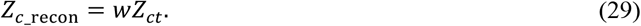

The residual representation was then defined as:

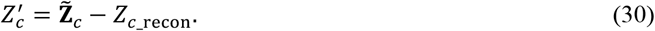

Here, 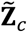 represents L2-normalized *Z_c_* and *Z_c_*__recon_ represents the component of *Z_c_* reconstructed from the estimated cell-type composition and reference cell-type prototypes, whereas *Z^’^_c_* represents the remaining information in *Z_c_* that is not explained by this reconstruction. *Z^’^_c_* should therefore be interpreted as a reconstruction residual rather than as a component completely independent of cell-type composition.

### Datasets and data preprocessing

DECIPHER was evaluated across transcriptomic, proteomic, metabolomic, spatial-transcriptomic, and regulatory-genomic settings. For benchmark datasets shared with previous deconvolution studies, the corresponding reference-target partitions were followed where applicable. Additional analyses included experimentally generated bulk-cell mixtures, immune-age prediction, and lung-cancer prognostic modeling.

### Real-bulk PBMC benchmark

For the real-bulk PBMC benchmark, we adopted the simulated training dataset released with TAPE^14^. The simulated samples contained profiles corresponding to five major immune cell populations (B cells, CD4^+^ T cells, CD8^+^ T cells, NK cells, and monocytes), with 2,000 shared highly variable genes. The 32,032 simulated tissue profiles provided with TAPE were used as supervised training data. The real target datasets comprised Monaco^35^ (n = 12), Microarray^10^ (n = 20), and SDY67^36^ (n = 12) from the corresponding published literature, for a total of 44 bulk-cell samples.

### CITE-seq RNA

The CITE-seq RNA dataset^32^ contained 11,012 cells from five immune cell populations and 1,970 shared genes. Donors HS1, HS3, and HS5 were used as the reference set to generate 3,000 simulated training mixtures, whereas HS12, HS13, and HS15 were held out to generate 3,000 independent simulated test mixtures.

### CITE-seq ADT

The CITE-seq ADT dataset^32^ used the same donor split as the RNA analysis above and contained 11,647 cells and 204 protein features. Three thousand simulated mixtures were generated for both the reference-derived training set and the held-out test set.

### Breast-tumor transcriptomics

The breast-tumor transcriptomic dataset^29^ contained 11,820 cells from six major populations: T cells, endothelial cells, cancer-associated fibroblasts, perivascular-like cells, myeloid cells, and B cells. CID4471, CID3586, and CID4067 were used as reference samples, whereas CID3838, CID4465, and CID4515 were used as independent test samples. Two thousand highly variable genes were retained.

### Pancreatic-islet transcriptomics

The pancreatic-islet dataset^30^ contained 17,134 cells from beta, alpha, delta, ductal, gamma, and acinar cell populations. VSG_MUC13639, VSG_MUC13640, and VSG_MUC13641 were used as reference samples, and VSG_MUC13631, VSG_MUC13632, and VSG_MUC13642 were used as independent test samples. Two thousand highly variable genes across the cell-types were retained.

### Bone-marrow metabolomics

The bone-marrow metabolomics dataset^33^ contained 652 single cells from five populations and 107 mass-spectrometry features. Cells were divided into reference and test subsets while preserving cell-type composition, and 1,000 simulated mixtures were generated from each subset.

### Breast proteomics

The breast proteomic dataset^34^ contained 59,255 cells from six major populations measured across 34 CyTOF protein markers. B1H09, B1H32, and B1H43 were used as reference samples, whereas B1H19, B1H24, and B1H49 were used as test samples; 3,000 simulated mixtures were generated independently from each predefined partition.

### Slide-seqV2 spatial transcriptomics

The Slide-seqV2 dataset^37^ contained 41,786 spatially resolved cells/beads assigned to 14 cell populations. Ungridded profiles were used to generate 6,000 simulated training mixtures. The original spatial data were further aggregated into 100 × 100 μm grids, yielding 1,892 target mixtures. Four thousand highly variable genes were retained for this analysis.

### Regulatory-genomics cross-modality datasets

For cross-modality regulatory-genomics evaluation, we followed the DeepDETAILS-style design^38^ consisting of six mixtures: transcription-initiation datasets 5D1 and 5D2, pause-release datasets 5D3 and 5D4, and histone-modification datasets 5D5 (H3K4me3) and 5D6 (H3K27ac). Each dataset contained five cell lineages, and 6,000 simulated training profiles were generated from the corresponding single-cell chromatin-accessibility reference.

First, mixed target-modality bulk profiles were encoded by DECIPHER into the domain-constant latent space *Z_c_*, while cell-type latent prototypes derived from the snATAC-seq reference were projected into the same latent space. Lineage proportions *π* were then estimated using differentiable non-negative least squares (NNLS), with the snATAC-seq-derived latent prototypes serving as the reference.

Second, with the estimated lineage proportions *π* fixed, non-negative cell-type-specific target-modality profiles were optimized to reconstruct the observed bulk signals, with a weak prior derived from the snATAC-seq reference. These profiles were combined with the estimated proportions to obtain cell-type-specific contributions and were subsequently rescaled to match the total bulk signal. Thus, this benchmark evaluates the recovery of cell-type-specific regulatory signals rather than cell-type proportion estimation alone.

### Immune-age PBMC datasets

For immune-age analysis, we used our previously published PBMC single-cell transcriptomic reference^60^ containing 361,911 cells and 449 healthy PBMC bulk-cell RNA expression samples with chronological age information spanning approximately 1-99 years. More than 30 original cell annotations were consolidated into 11 major groups: B cells, plasma cells, CD8^+^ naive T cells, CD8^+^ effector-memory T cells, CD8^+^ cytotoxic T cells, CD4^+^ memory T cells, monocytes, plasmacytoid dendritic cells, conventional dendritic cells, NK cells, and an “other cell types” group. After matching the single-cell and bulk datasets, 1,295 genes were retained and 6,000 simulated bulk profiles were generated.

### Lung-cancer datasets

For lung-cancer analysis, GSE131907^61^ was used as the single-cell reference. Fifteen tumor-microenvironment cell populations were retained from the existing annotations: Epithelial, Malignant, Fibroblasts, Endothelial, Monocytes, alveolar macrophages (Alveolar Mac), monocyte-derived macrophages (mo-Mac), Dendritic cells, MAST, CD4^+^ T, Treg, NK, B cells, non-Exhausted CD8^+^ T, and Exhausted CD8^+^ T. Bulk datasets included TCGA-LUAD^43^, TCGA-LUSC^62^, GSE81089^46^, and GSE143486^47^. 2,000 highly variable genes were retained after feature matching.

### Experimental real-bulk cell-line benchmark

The experimentally generated real-bulk benchmark consisted of ten human cell lines: THP1, A2780, SU-DHL-4, SU-DHL-10, NCI-H23, NCI-H2170, iCell-0144a, J76, HEK293T, and K562. The matched single-cell reference contained 11,319 cells after processing. A total of 18,451 genes were shared between the single-cell reference and bulk RNA-seq datasets, from which 2,000 highly variable genes were retained. Six thousand simulated bulk-cell profiles were generated for supervised DECIPHER training.

### Data preprocessing

Single-cell transcriptomic data were processed using Scanpy^63^. Genes detected in fewer than three cells were removed. Cells with fewer than 200 detected genes were excluded, and cells with mitochondrial gene expression fractions greater than or equal to 20% were additionally removed. Raw single-cell counts were normalized to 10,000 counts per cell followed by log1p transformation. For each dataset, genes shared between the single-cell reference and the corresponding target dataset were identified first; highly variable genes were then selected within this shared feature space using the single-cell reference and the Seurat method^64^. Most transcriptomic analyses retained 2,000 highly variable genes, whereas 4,000 genes were used for Slide-seqV2.

Simulated bulk profiles were generated by sampling cell-type proportions from a Uniform distribution and aggregating single-cell profiles according to the sampled compositions. Each simulated mixture was generated by aggregating exactly 200 cells. Raw count-based target data (real bulk omics) were transformed using counts per million (CPM) followed by log1p transformation. After feature matching, simulated and target profiles were normalized independently by their sample-wise maximum value. Proteomic and metabolomic datasets were processed using the same feature-matching and sample-wise normalization strategy.

## Data and code availability

All public datasets used in this study are publicly accessible through the original publications and repositories, with detailed dataset information and accession numbers provided in Supplementary Table 1. The experimentally generated bulk-cell RNA-seq dataset and matched single-cell RNA-seq reference dataset have been deposited in the NCBI Gene Expression Omnibus (GEO) under accession numbers GSE345995 and GSE345996, respectively. Python source code and DECIPHER are publicly available on GitHub at https://github.com/aapupu/DECIPHER. An online implementation of DECIPHER is available at https://luo-sysbiomed.cn/models/DECIPHER.

## Supplementary materials

Supplementary information includes Supplementary Figures S1-S4, Supplementary Table 1 (summary of datasets used in this study), Supplementary Table 2 (cell lines used in the mixture experiment), and Supplementary Table 3 (detailed methods comparison results). Supporting model files and benchmark results are available via Figshare at https://doi.org/10.6084/m9.figshare.33978271.

## Acknowledgements

This study was supported by funding from the National Key Research and Development Program of China (2025YFA1805302), the Natural Science Foundation of China (92370107 and T2541032), the Science and Technology Project in Guangzhou (202102070001), and the Fundamental Research Funds for the Central Universities (21626401). We thank Prof. Guobing Chen and members of his laboratory for kindly providing the THP1, A2780, J76, and HEK293T cell lines. O.J.L. acknowledges the support of the K. C. Wong Education Foundation and the Bingzheng Young Scholar Program of Jinan University.

## Author contributions

W.L. and O.J.L. conceived the study. W.L. and C.Li curated and processed the data. W.L. and C.Li implemented the model and conducted the computational experiments. Q.D., Y.Z. and Z.L. conducted the cell culturing and experiments for the cell-line mixture sequencing. W.L. and C.Li analyzed the results. C.Liu set up the web server. C.Li, W.L. and O.J.L. drafted the manuscript with input from the other authors.

## Competing interests

The authors declare no competing interests.

## Supplemental figure legends

**Figure S1:**
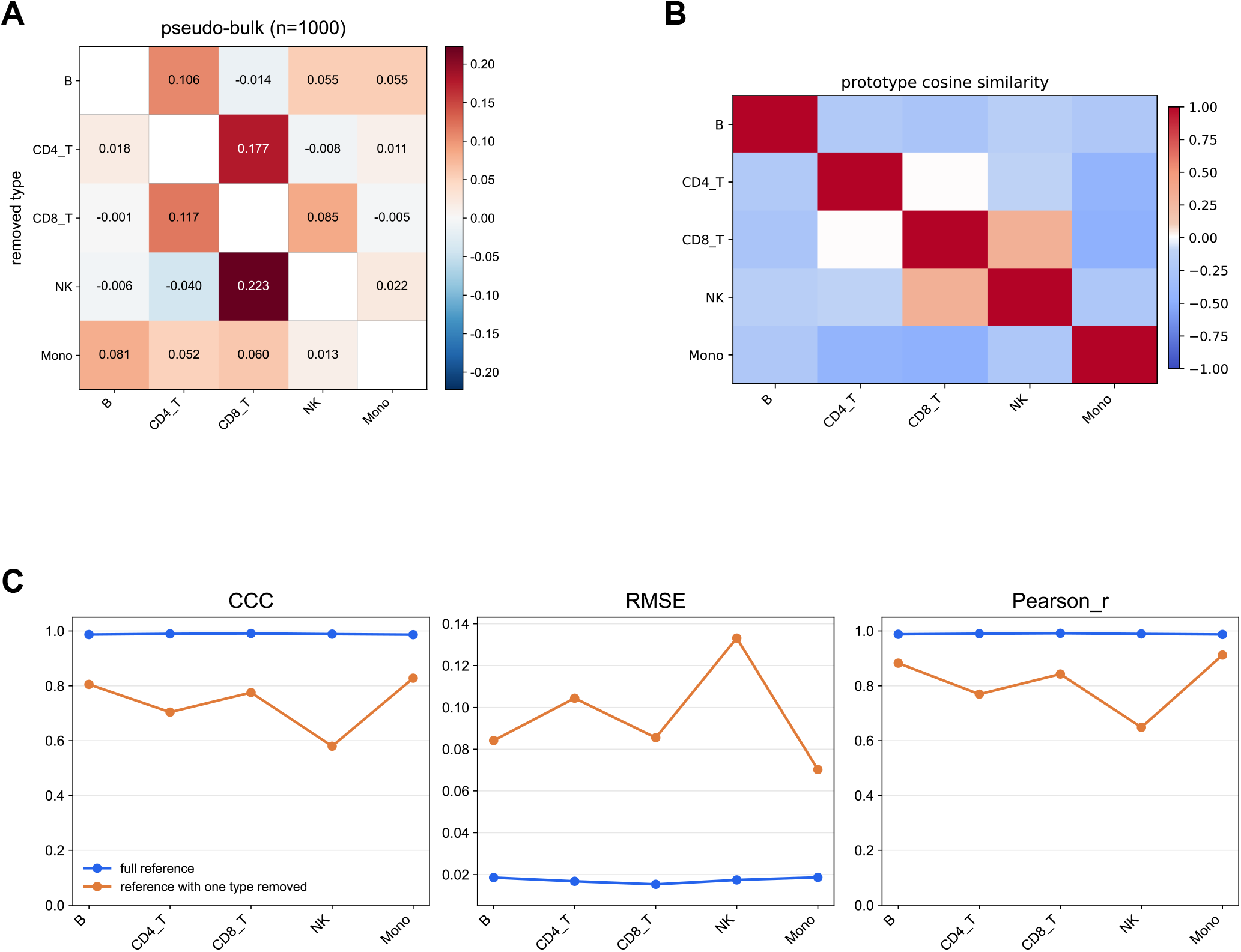
Effects of incomplete cell-type references on DECIPHER deconvolution. (A) Redistribution of predicted proportions among the remaining cell types after individually removing one cell type from the reference. Rows indicate the removed cell type and columns indicate the remaining cell types; red and blue denote increases and decreases, respectively, relative to the full-reference setting. (B) Cosine similarity among the cell-type latent prototypes (*Z*_ct_), with color indicating the magnitude and direction of similarity. (C) Changes in CCC, RMSE, and Pearson’s correlation coefficient (r) after individually removing one cell type from the reference. Blue lines indicate the full-reference setting, whereas orange lines indicate results after removal of the corresponding cell type.

**Figure S2:**
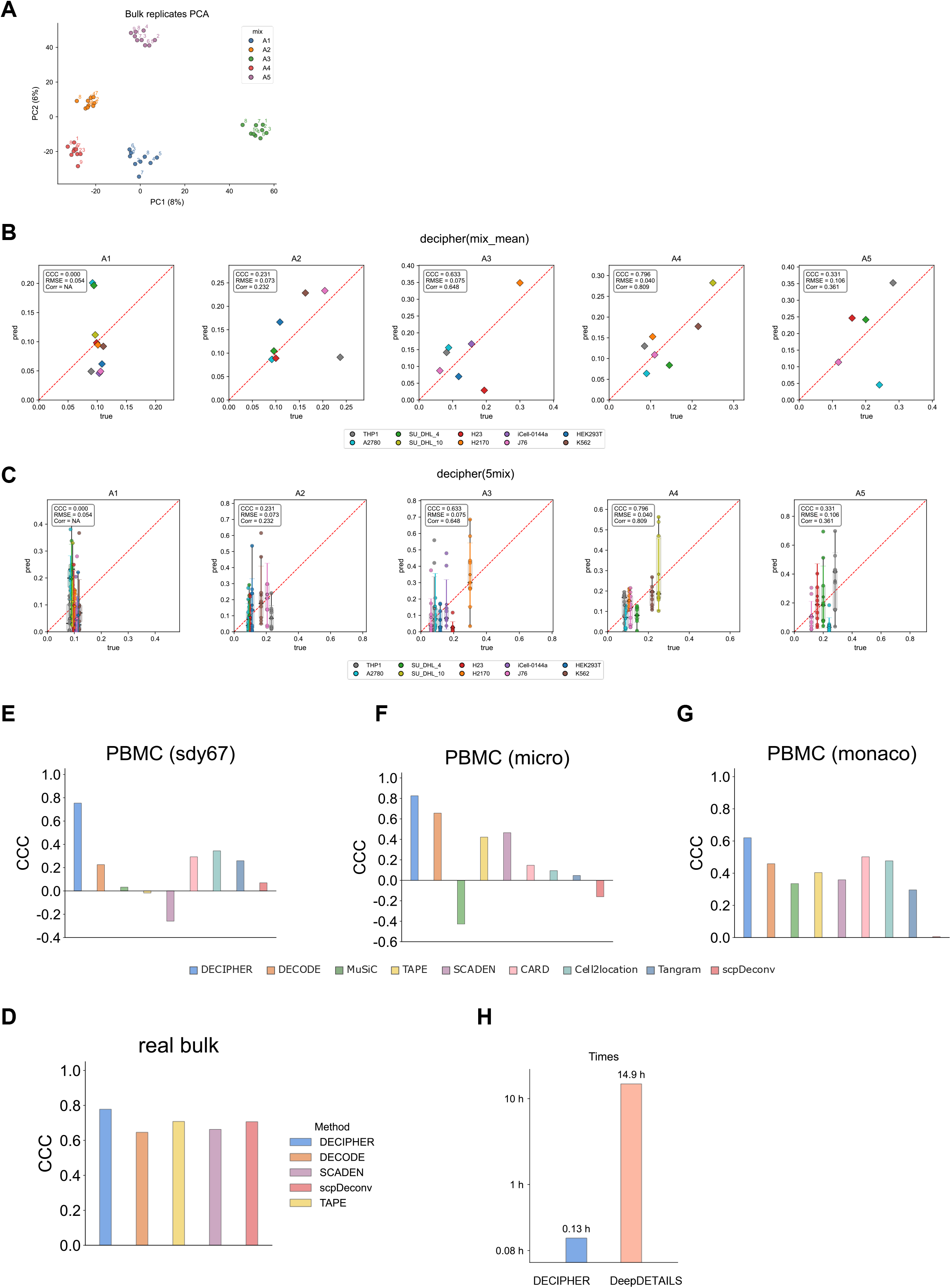
Characterization of experimentally generated bulk-cell mixture RNA-seq data and additional evaluation of DECIPHER deconvolution performance. (A) PCA plot of RNA-seq replicates of the five predefined cell-mixture designs (A1–A5). Each mixture was divided into 9 replicates for sequencing. (B) Cell-type proportion deconvolution results from DECIPHER across the five mixture designs (A1–A5). Mean predicted proportions across all bulk replicates within each mixture design are shown. The x-axis represents the true cell-type proportions, and the y-axis represents the predicted proportions. Colors denote different cell types, and the red dashed line indicates perfect agreement between predicted and true proportions. CCC, RMSE, and Pearson’s correlation coefficient (r) are shown for each mixture design. (C) Replicate-level deconvolution results across the five mixture designs (A1– A5), showing individual predicted cell-type proportions and their distributions for each bulk replicate. Axes, colors, and performance metrics are defined as in (B). (D) Overall deconvolution performance of DECIPHER and other deep-learning-based deconvolution methods on the experimentally generated real bulk-cell mixtures, evaluated using the same pooled-vector strategy. (E–G) Overall deconvolution performance of different methods on the SDY67 (E), Microarray (F), and Monaco (G) PBMC datasets, respectively. Because each dataset contains relatively few samples and limited variation in cell-type proportions, ground-truth and predicted proportions across all samples and cell types were pooled into two corresponding vectors for overall CCC calculation. (H) Comparison of the computational runtime of DECIPHER and DeepDETAILS for the same cross-modality deconvolution tasks.

**Figure S3:**
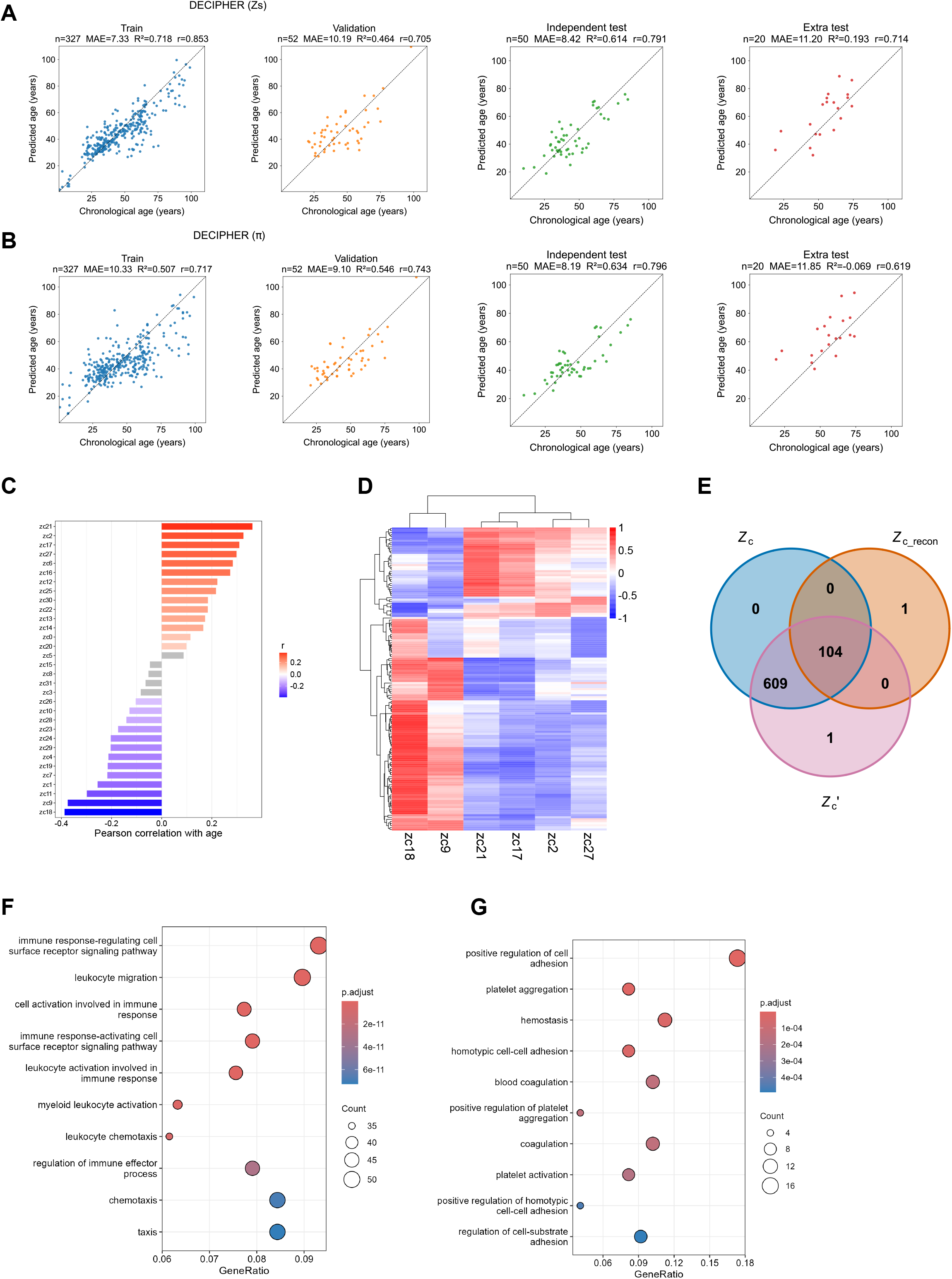
Age-associated analyses of DECIPHER representations and cell-type proportions. (A) Age prediction performance based on the DECIPHER domain-specific latent representation *Z*_s_. Scatter plots show the relationship between predicted age and chronological age across the training, validation, independent test, and additional test datasets. The sample size (n), mean absolute error (MAE), coefficient of determination (R²), and Pearson’s correlation coefficient (r) are indicated for each dataset. (B) Age-prediction performance using the DECIPHER-estimated cell-type proportions (π) as input features, shown using the same datasets and evaluation metrics as in (A). (C) Correlation analysis between the DECIPHER domain-constant latent representation *Z*_c_ and age. The bar plot shows Pearson’s correlation coefficient (r) between individual *Z*_c_ features and chronological age. Red and blue indicate positively and negatively age-associated features, respectively, whereas gray indicates features without significant correlations. (D) Biological interpretation of age-associated *Z*_c_ features. The heatmap displays Pearson’s correlation coefficient (r) between the top six age-associated *Z*_c_ features and their associated genes. Hierarchical clustering reveals coordinated patterns between latent features and gene expression profiles. (E) Venn diagram comparing gene sets associated with three sample-level representations for the same top six age-associated *Z*_c_ dimensions selected in (D): the full representation *Z*_c_, the composition-reconstructed representation *Z*_c_recon_ = *wZ*_ct_, and the reconstruction residual *Z*_c_^’^ = Z° _c_ − *Z*_c_recon_. *Z*_c_recon_ represents the component reconstructed from estimated cell-type proportions and cell-type prototypes, whereas *Z*_c_^’^ represents information not explained by this reconstruction, without implying complete independence from cellular composition. (F) Gene Ontology (GO) Biological Process enrichment analysis of the 609 genes shared by *Z*_c_ and *Z*_c_^’^ but not *Z*_c_recon_, representing the residual-associated gene set. The x-axis indicates GeneRatio, dot size represents gene count, and color denotes the adjusted P value. (G) Gene Ontology (GO) Biological Process enrichment analysis of the 104 genes shared by *Z*_c_, *Z*_c_recon_, and *Z*_c_^’^. The x-axis indicates GeneRatio, dot size represents gene count, and color denotes the adjusted P value.

**Figure S4:**
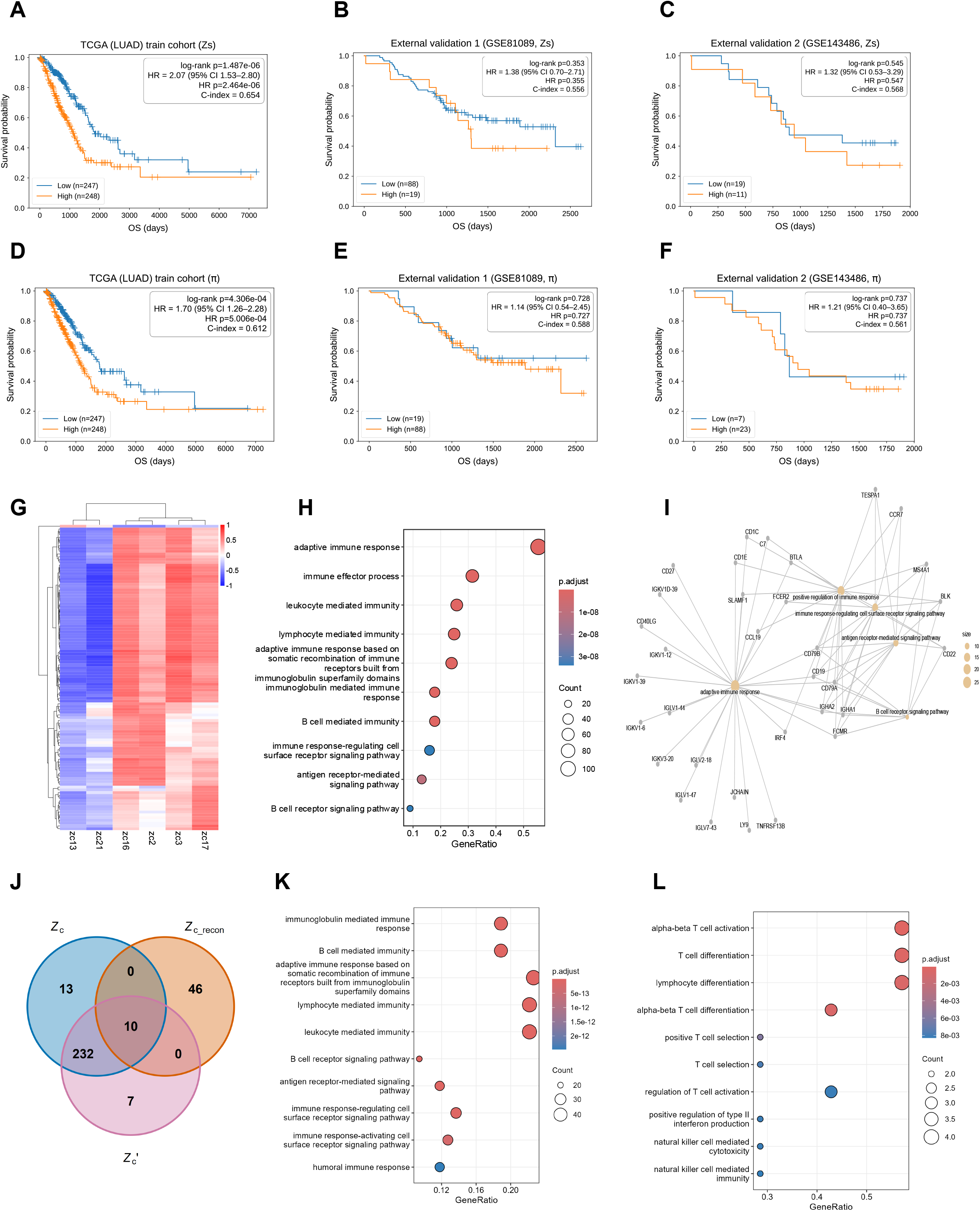
Prognostic analysis and biological interpretation of DECIPHER representations and cell-type proportions. (A–C) Kaplan–Meier overall survival curves for the high- and low-risk groups defined using risk scores derived from *Z*_s_ in the TCGA-LUAD training cohort (A), GSE81089 external validation cohort (B), and GSE143486 external validation cohort (C). Follow-up time in days is plotted against overall survival probability. Tick marks indicate censored observations, and n denotes the number of patients in each risk group. The log-rank P value assesses the difference in survival distributions between the two groups. HR denotes the hazard of death in the high-risk group relative to the low-risk group and is reported with its 95% confidence interval (CI). The concordance index (C-index) quantifies the ability of the model to discriminate between patients with different survival risks; values closer to 1 indicate better discrimination, whereas a value of 0.5 indicates performance comparable to random prediction. (D–F) Kaplan–Meier overall survival curves based on risk scores derived from the DECIPHER-estimated cell-type proportions (π) in the TCGA-LUAD training cohort (D), GSE81089 external validation cohort (E), and GSE143486 external validation cohort (F), using the same survival metrics as in (A–C). (G) Correlation heatmap between the six *Z*_c_ features most significantly associated with survival identified in the TCGA-LUAD training cohort and their top 30 associated genes after deduplication. The heatmap shows Pearson’s correlation coefficient (r) between *Z*_c_ features and gene expression profiles in lung adenocarcinoma samples. Hierarchical clustering reveals correlation patterns among genes and latent features, with colors indicating the magnitude and direction of correlations. (H) Gene Ontology (GO) Biological Process enrichment analysis of genes positively correlated with the prognostic latent feature *Z*_c16_. *Z*_c16_ was selected as the latent feature with the largest absolute Cox regression coefficient among the standardized *Z*_c_ features in the TCGA-LUAD training cohort. In the dot plot, the x-axis indicates the GeneRatio, dot size represents the number of genes associated with each GO term, and color denotes the adjusted P value. (I) Gene–GO term network showing the relationships between enriched GO terms and their associated genes identified in (H). (J) Venn diagram comparing gene sets associated with the full representation *Z*_c_, the composition-reconstructed representation *Z*_c_recon_, and the reconstruction residual *Z*_c_^’^ for the same six *Z*_c_ dimensions associated with prognosis selected in (G). (K) Gene Ontology (GO) Biological Process enrichment analysis of the 240 genes shared by *Z*_c_ and *Z*_c_^’^ but not *Z*_c_recon_. The x-axis indicates GeneRatio, dot size represents gene count, and color denotes the adjusted P value. (L) Gene Ontology (GO) Biological Process enrichment analysis of the 10 genes shared by *Z*_c_, *Z*_c_recon_, and *Z*_c_^’^. The x-axis indicates GeneRatio, dot size represents gene count, and color denotes the adjusted P value.

## Notes

https://github.com/aapupu/DECIPHER

https://luo-sysbiomed.cn/models/DECIPHER

https://doi.org/10.6084/m9.figshare.33978271

### Competing Interest Statement

The authors have declared no competing interest.

### Summary of Updates

Spelling, grammar and formatting corrections. Duplicated cell line names were corrected.

## References

1 GTEx Consortium. The GTEx Consortium atlas of genetic regulatory effects across human tissues. Science 369, 1318–1330 (2020).

2 Liu, J., Lichtenberg, T. & Hoadley, K. A. An Integrated TCGA Pan-Cancer Clinical Data Resource to Drive High-Quality Survival Outcome Analytics. Cell 173, 400–416.e411 (2018).

3 Garmire, L. X., Li, Y. & Huang, Q. Challenges and perspectives in computational deconvolution of genomics data. Nat. Methods 21, 391–400 (2024).

4 Avila Cobos, F., Alquicira-Hernandez, J., Powell, J. E., Mestdagh, P. & De Preter, K. Benchmarking of cell type deconvolution pipelines for transcriptomics data. Nat. Commun. 11, 5650 (2020).

5 Tsoucas, D. et al. Accurate estimation of cell-type composition from gene expression data. Nat. Commun. 10, 2975 (2019).

6 Regev, A. et al. The Human Cell Atlas. eLife 6, e27041 (2017).

7 Tabula Sapiens Consortium. The Tabula Sapiens: A multiple-organ, single-cell transcriptomic atlas of humans. Science 376, eabl4896 (2022).

8 Wang, X., Park, J., Susztak, K., Zhang, N. R. & Li, M. Bulk tissue cell type deconvolution with multi-subject single-cell expression reference. Nat. Commun. 10, 380 (2019).

9 Newman, A. M., Steen, C. B. & Liu, C. L. Determining cell type abundance and expression from bulk tissues with digital cytometry. Nat. Biotechnol. 37, 773–782 (2019).

10 Newman, A. M., Liu, C. L. & Green, M. R. Robust enumeration of cell subsets from tissue expression profiles. Nat. Methods 12, 453–457 (2015).

11 Erdmann-Pham, D. D., Fischer, J., Hong, J. & Song, Y. S. Likelihood-based deconvolution of bulk gene expression data using single-cell references. Genome Res. 31, 1794–1806 (2021).

12 Andrade Barbosa, B., et al. Bayesian log-normal deconvolution for enhanced in silico microdissection of bulk gene expression data. Nat. Commun. 12, 6106 (2021).

13 Menden, K., Marouf, M. & Oller, S. Deep learning-based cell composition analysis from tissue expression profiles. Sci. Adv. 6, eaba2619 (2020).

14 Chen, Y., Wang, Y. & Chen, Y. Deep autoencoder for interpretable tissue-adaptive deconvolution and cell-type-specific gene analysis. Nat. Commun. 13, 6735 (2022).

15 Zhao, T., Liu, R. & Sun, Y. DECODE: deep learning-based common deconvolution framework for various omics data. Nat. Methods 23, 596–608 (2026).

16 Wang, F., Yang, F. & Huang, L. Deep domain adversarial neural network for the deconvolution of cell type mixtures in tissue proteome profiling. *Nat*. Mach. Intell. 5, 1236– 1249 (2023).

17 Kleshchevnikov, V., Shmatko, A. & Dann, E. Cell2location maps fine-grained cell types in spatial transcriptomics. Nat. Biotechnol. 40, 661–671 (2022).

18 Biancalani, T., Scalia, G. & Buffoni, L. Deep learning and alignment of spatially resolved single-cell transcriptomes with Tangram. Nat. Methods 18, 1352–1362 (2021).

19 Ma, Y. & Zhou, X. Spatially informed cell-type deconvolution for spatial transcriptomics. Nat. Biotechnol. 40, 1349–1359 (2022).

20 Bengio, Y., Courville, A. & Vincent, P. Representation Learning: A Review and New Perspectives. IEEE Transactions on Pattern Analysis and Machine Intelligence 35, 1798– 1828 (2013).

21 Li, W. et al. Single-cell immune aging clocks reveal inter-individual heterogeneity during infection and vaccination. *Nat*. Aging 5, 607–621 (2025).

22 Chu, T., Wang, Z., Pe’er, D. & Danko, C. G. Cell type and gene expression deconvolution with BayesPrism enables Bayesian integrative analysis across bulk and single-cell RNA sequencing in oncology. *Nat*. Cancer 3, 505–517 (2022).

23 Theodoris, C. V., Xiao, L. & Chopra, A. Transfer learning enables predictions in network biology. Nature 618, 616–624 (2023).

24 Cui, H., Wang, C. & Maan, H. scGPT: toward building a foundation model for single-cell multi-omics using generative AI. Nat. Methods 21, 1470–1480 (2024).

25 Jew, B., Alvarez, M. & Rahmani, E. Accurate estimation of cell composition in bulk expression through robust integration of single-cell information. Nat. Commun. 11, 1971 (2020).

26 Luecken, M. D., Büttner, M. & Chaichoompu, K. Benchmarking atlas-level data integration in single-cell genomics. Nat. Methods 19, 41–50 (2022).

27 Goh, W. W. B., Wang, W. & Wong, L. Why Batch Effects Matter in Omics Data, and How to Avoid Them. Trends Biotechnol. 35, 498–507 (2017).

28 Lin, L. I. A concordance correlation coefficient to evaluate reproducibility. Biometrics 45, 255–268 (1989).

29 Wu, S. Z., Al-Eryani, G. & Roden, D. L. A single-cell and spatially resolved atlas of human breast cancers. Nat. Genet. 53, 1334–1347 (2021).

30 Oppenländer, L. Vertical sleeve gastrectomy triggers fast β-cell recovery upon overt diabetes. Mol. Metab. 54, 101330 (2021).

31 Stoeckius, M. et al. Simultaneous epitope and transcriptome measurement in single cells. Nat. Methods 14, 865–868 (2017).

32 He, J. Y. Dysregulation of CD4+ and CD8+ resident memory T, myeloid, and stromal cells in steroid-experienced, checkpoint inhibitor colitis. J. Immunother. Cancer 12, e008628 (2024).

33 Cao, J. Deciphering the metabolic heterogeneity of hematopoietic stem cells with single-cell resolution. Cell Metab. 36, 209–221.e206 (2024).

34 Gray, G. K. A human breast atlas integrating single-cell proteomics and transcriptomics. Dev. Cell 57, 1400–1420.e1407 (2022).

35 Monaco, G., Lee, B. & Xu, W. RNA-Seq signatures normalized by mRNA abundance allow absolute deconvolution of human immune cell types. Cell Rep. 26, 1627–1640.e1627 (2019).

36 Zimmermann, M. T. System-wide associations between DNA-methylation, gene expression, and humoral immune response to influenza vaccination. PLoS One 11, e0152034 (2016).

37 Stickels, R. R., Murray, E. & Kumar, P. Highly sensitive spatial transcriptomics at near-cellular resolution with Slide-seqV2. Nat. Biotechnol. 39, 313–319 (2021).

38 Yao, L., Shah, S. R. & Ozer, A. High-resolution reconstruction of cell-type-specific transcriptional regulatory processes from bulk sequencing samples. Nat. Biotechnol., (2026).

39 Becht, E., McInnes, L. & Healy, J. Dimensionality reduction for visualizing single-cell data using UMAP. Nat. Biotechnol. 37, 38–44 (2019).

40 Gene Ontology Consortium. The Gene Ontology knowledgebase in 2023. Genetics 224, iyad031 (2023).

41 Blackburn, S. D. et al. Coregulation of CD8+ T cell exhaustion by multiple inhibitory receptors during chronic viral infection. Nat. Immunol. 10, 29–37 (2009).

42 Quigley, M. et al. Transcriptional analysis of HIV-specific CD8+ T cells shows that PD-1 inhibits T cell function by upregulating BATF. Nat. Med. 16, 1147–1151 (2010).

43 Cancer Genome Atlas Research Network. Comprehensive molecular profiling of lung adenocarcinoma. Nature 511, 543–550 (2014).

44 Cox, D. R. Regression Models and Life-Tables. Journal of the Royal Statistical Society: Series B 34, 187–220 (1972).

45 Kaplan, E. L. & Meier, P. Nonparametric Estimation from Incomplete Observations. J. Am. Stat. Assoc. 53, 457–481 (1958).

46. Djureinovic, D., Hallström, B. M. & Horie, M. Profiling cancer testis antigens in non-small-cell lung cancer. JCI Insight 1, e86837 (2016).

47 Zhang, Z. Reference genome and annotation updates lead to contradictory prognostic predictions in gene expression signatures: a case study of resected stage I lung adenocarcinoma. Brief. Bioinform. 22, bbaa081 (2021).

48 Lawson, C. L. & Hanson, R. J. Solving Least Squares Problems. (Society for Industrial and Applied Mathematics, 1995).

49 Kedzierska, K. Z., Crawford, L., Amini, A. P. & Lu, A. X. Zero-shot evaluation reveals limitations of single-cell foundation models. Genome Biol. 26, (2025).

50 Beck, A. & Teboulle, M. Mirror descent and nonlinear projected subgradient methods for convex optimization. Oper. Res. Lett. 31, 167–175 (2003).

51 Gretton, A., Borgwardt, K. M., Rasch, M. J., Schölkopf, B. & Smola, A. A Kernel Two-Sample Test. J. Mach. Learn. Res. 13, 723–773 (2012).

52 Loshchilov, I. & Hutter, F. in International Conference on Learning Representations (ICLR 2019) (2019).

53 Paszke, A. et al. in Advances in Neural Information Processing Systems 32 (NeurIPS 2019) Vol. 32 8024–8035 (2019).

54 FastQC: a quality control tool for high throughput sequence data (Babraham Bioinformatics, 2010).

55 Trim Galore! (Babraham Bioinformatics, 2015).

56 Kim, D., Paggi, J. M., Park, C., Bennett, C. & Salzberg, S. L. Graph-based genome alignment and genotyping with HISAT2 and HISAT-genotype. Nat. Biotechnol. 37, 907–915 (2019).

57 Liao, Y., Smyth, G. K. & Shi, W. featureCounts: an efficient general purpose program for assigning sequence reads to genomic features. Bioinformatics 30, 923–930 (2014).

58 Zheng, G. X. Y., Terry, J. M. & Belgrader, P. Massively parallel digital transcriptional profiling of single cells. Nat. Commun. 8, 14049 (2017).

59 Wu, T. et al. clusterProfiler 4.0: A universal enrichment tool for interpreting omics data. The Innovation 2, (2021).

60 Lai, W., Feng, Q. & Lei, W. Deciphering Immunosenescence From Child to Frailty: Transcriptional Changes, Inflammation Dynamics, and Adaptive Immune Alterations. Aging Cell 24, e70082 (2025).

61 Kim, N., Kim, H. K. & Lee, K. Single-cell RNA sequencing demonstrates the molecular and cellular reprogramming of metastatic lung adenocarcinoma. Nat. Commun. 11, 2285 (2020).

62. Cancer Genome Atlas Research Network. Comprehensive genomic characterization of squamous cell lung cancers. Nature 489, 519–525 (2012).

63 Wolf, F. A., Angerer, P. & Theis, F. J. SCANPY: large-scale single-cell gene expression data analysis. Genome Biol. 19, 15 (2018).

64 Stuart, T., Butler, A. & Hoffman, P. Comprehensive Integration of Single-Cell Data. Cell 177, 1888–1902.e1821 (2019).

